# Molecular organization of soluble type III secretion system sorting platform complexes

**DOI:** 10.1101/648071

**Authors:** Ivonne Bernal, Jonathan Börnicke, Johannes Heidemann, Dmitri Svergun, Anne Tuukkanen, Charlotte Uetrecht, Michael Kolbe

## Abstract

Many medically relevant Gram-negative bacteria use the type III secretion system (T3SS) to translocate effector proteins into the host for their invasion and intracellular survival. A multi-protein complex located at the cytosolic interface of the T3SS is proposed to act as a sorting platform by selecting and targeting substrates for secretion through the system. However, the precise stoichiometry and 3D organization of the sorting platform components is unknown. Here we reconstitute soluble complexes of the *Salmonella* Typhimurium sorting platform proteins including the ATPase InvC, the regulator OrgB, the protein SpaO and a recently identified subunit SpaO_C_, which we show to be essential for the solubility of SpaO. We establish domain-domain interactions, determine for the first time the stoichiometry of each subunit within the complexes by native mass spectrometry and gain insight into their organization using small-angle X-ray scattering. Importantly, we find that in solution the assembly of SpaO/SpaO_C_/OrgB/InvC adopts an extended L-shaped conformation resembling the sorting platform pods seen in *in situ* cryo-electron tomography, proposing that this complex is the core building block that can be conceivably assembled into higher oligomers to form the T3SS sorting platform. The determined molecular arrangements of the soluble complexes of the sorting platform provide important insights into its architecture and assembly.

## Introduction

Type III secretion systems (T3SS) are protein nanomachines used by several medically relevant pathogenic Gram-negative bacteria to deliver effector molecules into host cells to subvert multiple cellular processes, leading to diseases such as salmonellosis, bubonic plague or sexually transmitted infections (1,2). The T3SS forms a syringe-shaped macromolecular complex of ~3.5 MDa, whose main elements are a basal body that spans both bacterial membranes and a protruding needle that forms a continuous secretion channel connecting the bacterial and host cell cytoplasms (3–5). The precise assembly and function of the T3SS critically depends on the hierarchical delivery of structural proteins to build the extracellular needle, followed by effector proteins for translocation into the host cell (6,7). The control of this ordered process involves a multi-protein complex associated with the cytoplasmic side of the T3SS that is proposed to act as a sorting platform by recognizing and selecting substrates for secretion through the system (8,9).

In *Salmonella* Typhimurium, the components of the sorting platform of the SPI-1 (Salmonella pathogenicity island 1) T3SS include the ATPase InvC (SctN in unified nomenclature), the protein SpaO (SctQ), the ATPase regulator OrgB (SctL) and the accessory protein OrgA (SctK) (8). Visualization of the SPI-1 sorting platform by cryo-electron tomographic analysis (CET) indicates that it adopts a structure of six pods containing SpaO that are connected to the T3SS base through OrgA and to a presumed hexameric ATPase through OrgB linkers (10,11). However, this contrasts with other studies on both the *S.* Typhimurium SPI-1 and the *Yersinia enterocolitica* T3SS showing the presence of ~24 and ~22 subunits, respectively, of the SctQ protein at the needle base, which suggests a more extensive structure comparable to the continuous cytosolic ring of flagellar T3SSs (12,13). Furthermore, the sorting platform has been found to be a dynamic structure in which different components are exchanging between a T3SS-associated state and a cytosolic pool (12–14).

The probably best characterized component in the *Salmonella* SP1-I sorting platform is the protein SpaO. SpaO contains two surface presentation of antigen domains (SPOA1 and SPOA2) that can form SPOA2-SPOA2 homodimers, as well as SPOA1-SPOA2 heterodimers that are able to interact with OrgB (15). Similar to the homologs of other pathogenic bacteria including *Yersinia* and *Shigella* species, the gene encoding SpaO contains an internal translation initiation site and produces an additional short isoform comprising the SPOA2 domain of SpaO, which we refer to as SpaO_C_ (16,17). This short product interacts with the full-length protein in other species and thus could represent an additional structural component of the sorting platform (12,18–20). However, the function of SpaO_C_ in type III secretion is elusive, and how it interacts with the other subunits of the sorting platform is unknown. Moreover, the precise protein composition and spatial molecular organization of the sorting platform, as well its assembly process and mechanism of action in substrate sorting remain uncertain.

In this study, we reconstitute and analyze for the first time the soluble assembling units of the *Salmonella* Typhimurium SPI-1 sorting platform using purified proteins. We observe that SpaO_C_, the second protein product of the gene *spaO*, is required for fully efficient type III function and for the stability of the sorting platform complexes in solution. Using native mass spectrometry (MS), small-angle X-ray scattering (SAXS) and multi-angle light scattering (MALS), we characterize different substructures of the sorting platform, determining their stoichiometry and association into SpaO/SpaO_C_/OrgB/InvC complexes. These complexes adopt an extended L-shaped conformation in solution that mirrors a segment of the sorting platform visualized by CET. Our data present the most detailed assembly of the *Salmonella* Typhimurium SPI-1 sorting platform in solution, reporting the conformation of what we propose is the core building block to assemble the sorting platform at the T3SS needle base.

## Results

### The *spaO* gene encodes two protein products required for fully active type III secretion

SpaO is a critical component of the *S.* Typhimurium SPI-1 sorting platform and it has recently been shown that the *spaO* gene, similar to several of its homologs in other T3SSs, produces both the full-length SpaO protein and a shorter variant that is the result of translation initiation from an internal ribosome binding site (RBS) (17). When we recombinantly expressed C-terminally *Strep*-tagged *spaO* and purified the protein by *Strep-*Tactin affinity purification, we could confirm the production of this smaller protein product SpaO_C_ (Fig. 1A). Using MALDI MS and Edman sequencing we found that it begins with a methionine, rather than a valine that is encoded at its starting position at codon 203 (Fig. S1 and Table S1), supporting the conclusion that it is the product of internal translation initiation.

**Figure 1.**
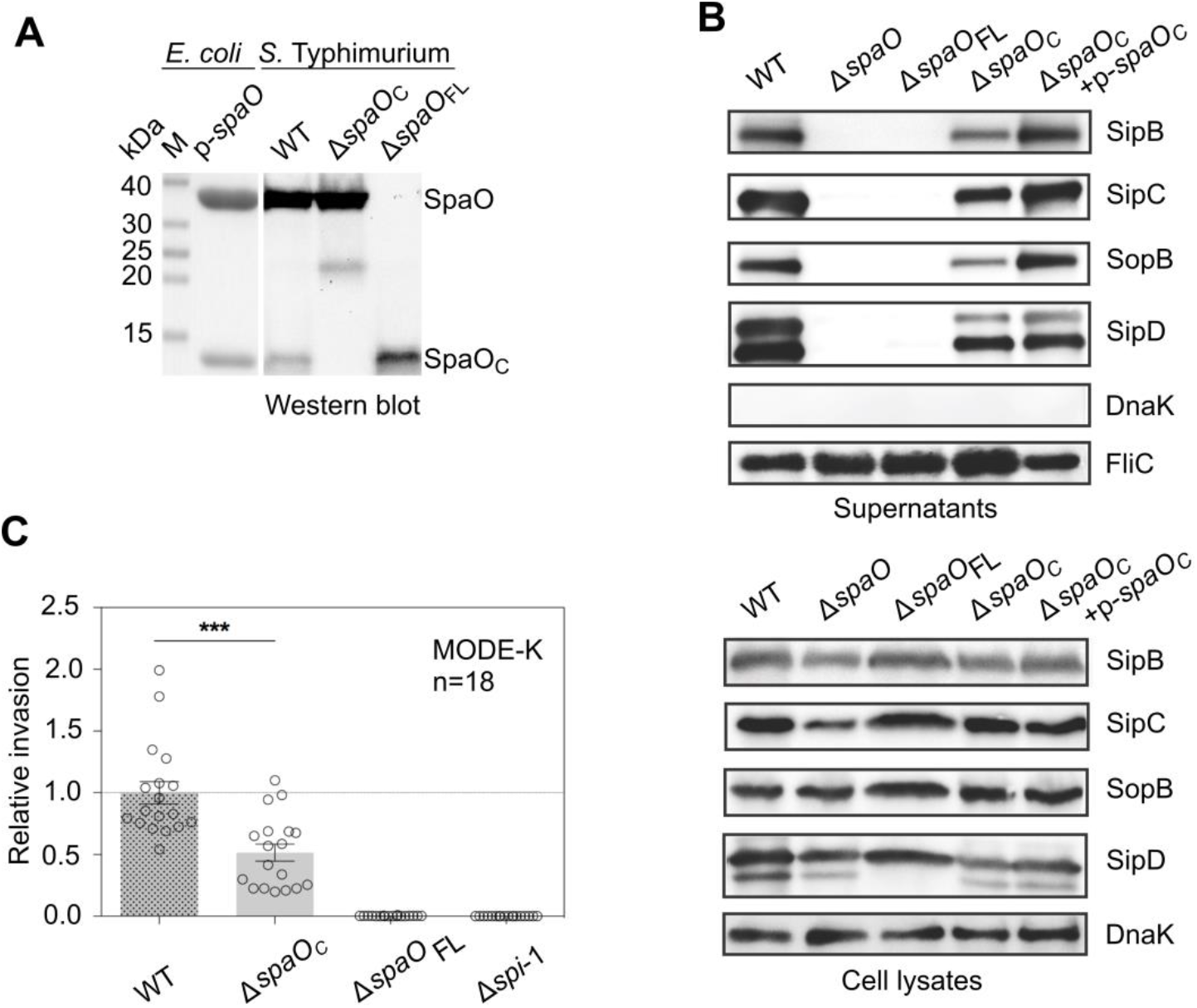
SpaO_C_ is made by internal translation initiation within the *spaO* gene and is required for fully efficient T3 secretion. (A) Coomassie-stained SDS-PAGE of *Strep*-tagged SpaO and SpaO_C_ purified from *E. coli* recombinantly expressing *spaO-Strep* (p-*spaO*) (left panel). Western blot detection of C-terminally 3xFLAG-tagged SpaO and SpaO_C_ in whole cell lysates of *Salmonella* wild type (WT) and strains harboring silent mutations at the *spaO* internal Shine-Dalgarno sequence and start codon (Δ*SpaO_C_*) or a nonsense mutation shortly after the *spaO* start codon (Δ*spaO_FL_*) (right panel). Molecular weight markers (M) are indicated and the result shown is representative of three biological replicates. (B) Western blot analysis of proteins secreted by *Salmonella* into culture supernatants (top panel). The proteins DnaK and FliC served as cell lysis control and loading control, respectively. Expression of T3SS substrates in whole cell lysates is shown in the bottom panel. Data shown are representative of three biological replicates. (C) Analysis of *Salmonella* invasion into MODE-K cells. Relative invasion was normalized to the levels of the wildtype and the results summarize three independent experiments. A *Salmonella* strain from which the entire *Salmonella* pathogenicity island 1 that encodes the SPI-1 T3SS has been deleted was included as a non-invasive control (Δ*spi-1*). Error bars represent one standard deviation. ***= p-value < 0.001

To examine the role of the SpaO isoforms in the infection process of *S.* Typhimurium, we first created mutants that produce only the full-length or short variant of SpaO by introducing into the chromosome either two stop codons shortly after the *spaO* start codon (Δ*spaO_FL_*) or silent mutations in both the putative RBS and start codon of SpaO_C_ (Δ*SpaO_C_*) (Fig. 1A). We tested the ability of these mutants to secrete T3SS substrate proteins into the culture supernatant, which showed that loss of spaO_FL_ causes complete inhibition of T3SS function similar to that observed for *spaO* gene knockout mutants (Fig. 1B). In contrast, abrogation of SpaO_C_ translation resulted in a marked reduction in secretion, which could almost completely be restored by complementation with *SpaO_C_*. Similarly, while the deletion of SpaO completely abolished the ability of *Salmonella* to invade host cells, loss of SpaO_C_ resulted in a statistically significant reduction of invasiveness by about 50% compared to the wild type (Fig. 1C). However, it is possible that the loss of SpaO_C_ in the *ΔSpaO_C_* mutant is incomplete, because when we expressed and affinity-purified a SpaO variant in which only the start codon of SpaO_C_ was mutated from GTG to GCG (SpaO_V203A_), small amounts of SpaO_C_ were still co-purified with the full-length SpaO (Fig. 2A). MALDI MS of this SpaO_C_ showed the presence of both methionine and alanine in the first amino acid position (Fig. S2, Table S2), indicating that even in the absence of internal translation initiation a SpaO_C_-like protein can still be produced by an alternative mechanism, probably proteolysis.

**Figure 2.**
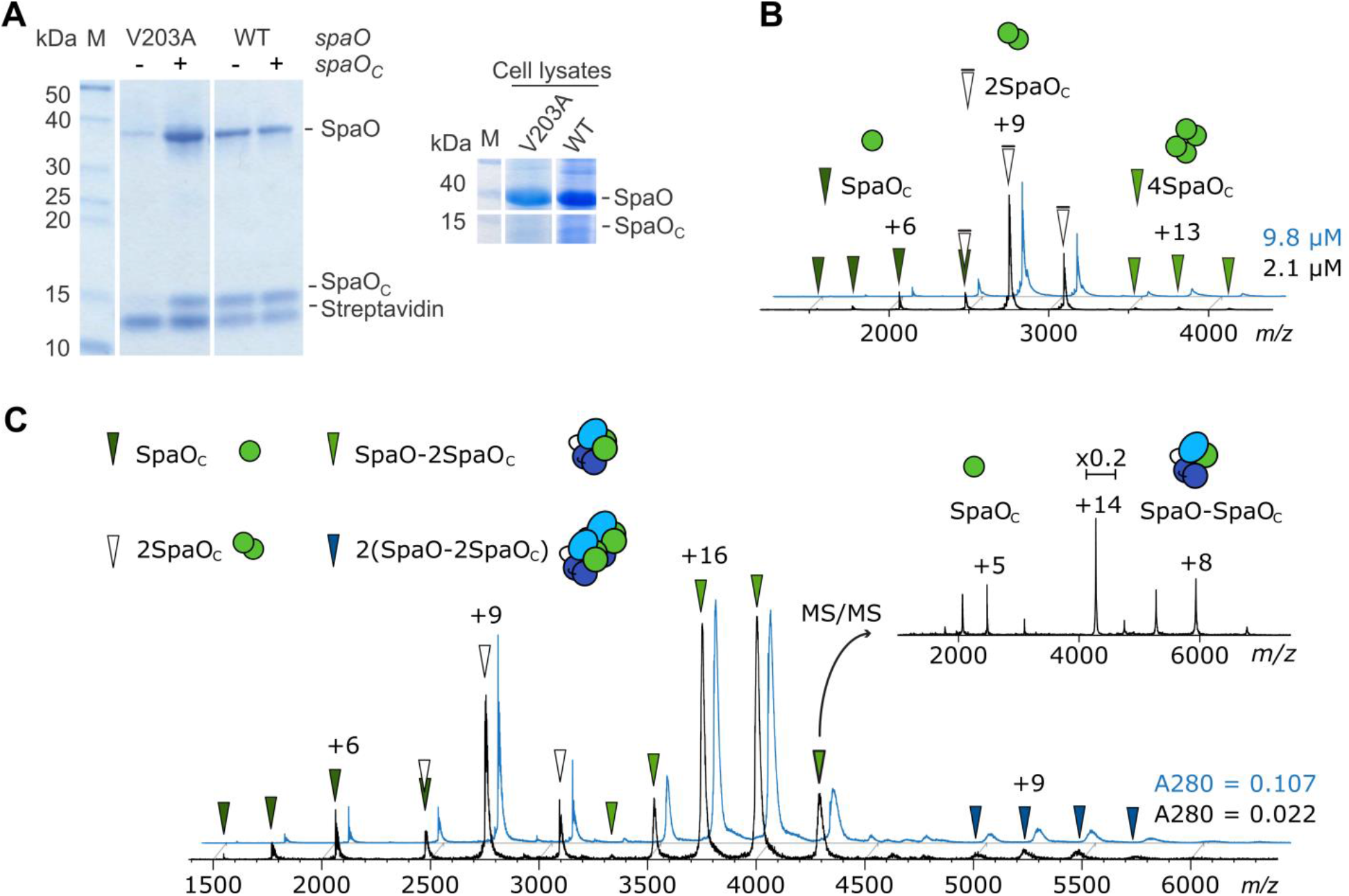
SpaO and SpaO_C_ interact to form stable 1:2 complexes. (A) Coomassie-stained SDS-PAGE of plasmid-encoded SpaO expressed in *Salmonella* Δ*spaO* and affinity-purified using *Strep*-Tactin. The SpaO V203A strain has a mutation at the SpaO_C_ start codon. The solubility of SpaO for the mutants was recovered by co-expression with plasmid-encoded *SpaO_C_* (+). A Coomassie-stained SDS-PAGE of whole cells lysates showing expression levels for SpaO and SpaO_C_ is depicted in the right panel. Molecular mass markers are included in lane M. (B) Native mass spectrum of SpaO_C_ at two different protein concentrations (black and blue spectra). SpaO_C_ monomers (dark green arrows), dimers (white arrows) and tetramers (light green arrows) are indicated. The main charge state of each protein or protein complex is labeled. (C) Native mass spectrum of SpaO/SpaO_C_ complexes at two different protein concentrations (black and blue spectra). The formation of SpaO-2SpaO_C_ complexes (light green arrows) and further dimerization of these heterotrimers (dark blue arrows) was observed irrespective of the protein concentration (indicated by absorbance at 280 nm, A280). CID MS/MS (inset) of the +14 precursor of the heterotrimer shows dissociation of SpaO_C_ monomers and a residual SpaO-SpaO_C_ complex. Both SpaO and SpaO_C_ carry a C-terminal *Strep*-tag. The precursor peak in the MS/MS spectrum has been scaled down to 20% of its original size. Experimental and theoretical molecular masses are given in Table S3.

### SpaO_C_ dimers bind to the N-terminal domain of SpaO to form SpaO-2SpaO_C_ complexes

In order to determine the molecular function of SpaO_C_ in type III secretion, we first tested the influence of SpaO_C_ on SpaO stability. To this end, we expressed C-terminally *Strep-*tagged SpaO_V203A_ in a *spaO* knockout strain and purified it using *Strep*-Tactin affinity chromatography. Interestingly, this mutation did not only almost completely abolish the production of soluble SpaO_C_, but also drastically reduced the levels of soluble full-length SpaO_V203A_ (Fig. 2A), which instead formed insoluble inclusion bodies (data not shown). Complementation of the mutant with *SpaO_C_* restored the soluble levels of both proteins, indicating that SpaO_C_ is required for SpaO stability in solution.

The observed enhancement of SpaO solubility could involve interaction between SpaO and SpaO_C_, as reported for homologous proteins (18–20). Therefore, we used size-exclusion chromatography (SEC) coupled to MALS and SAXS, as well as native MS to study complex formation between the two SpaO isoforms. In native MS non-covalent complexes of biological samples are ionized and transferred to the gas phase under mild conditions, making it a sensitive technique to take a snapshot of all non-covalent assemblies in a sample (21–23). First, we observed that SpaO_C_ exists mostly as a homodimer in solution (Fig. 2B and Fig. S3A, Table S4, SASDC68), consistent with the crystal structure of the SPOA2-SPOA2 domain dimer of SpaO and homolog proteins in *Shigella* and *Yersinia* (15,19,20). A very low abundance of homotetramers was observed irrespective of the protein concentration tested (Fig. 2B), indicating that these complexes reflect biologically relevant units and are not the result of unspecific clustering during the MS ionization process.

Subsequent analysis of co-purified SpaO/SpaO_C_ showed that both proteins interact to form predominantly heterotrimers with a stoichiometry of SpaO-2SpaO_C_ (Fig. 2C and Fig. S3B, Table S4, SASDC78). Notably, no monomeric SpaO was detected in MS measurements, which indicates high binding affinity within the SpaO-2SpaO_C_ complex and further highlights the critical role of SpaO_C_ in SpaO solubility. Excess SpaO_C_ was found to be mainly dimeric, suggesting that it binds to SpaO as a pre-formed dimer. In addition to the predominant SpaO-2SpaO_C_ species, we also observed the dimerization of these heterotrimers into 2(SpaO-2SpaO_C_) heterohexamers, which was independent of both the protein concentrations and the position of the *Strep*-tag (Fig. 2C, Fig. S3C). Higher-order oligomers could only be observed when measuring highly concentrated samples that showed unspecific clustering during the ionization process and were therefore considered to be non-specific assemblies. This conclusion is also supported by SEC-MALS, which at a high protein concentration of 140 μM showed no evidence of species larger than the 2(SpaO-2SpaO_C_) heterohexamer (Fig. S3B). Selected ions of the SpaO-2SpaO_C_ heterotrimers were subjected to collision-induced dissociation (CID) MS/MS experiments. The observed dissociation pathways in these experiments give further insights into complex topology since dissociating proteins are mostly small monomeric proteins from the periphery of protein complexes, although the process of CID is not completely understood and exceptions have been reported (24). Here, one SpaO_C_ monomer was found to dissociate, leaving a residual SpaO-SpaO_C_ complex (Fig. 2C inset).

In order to determine the domain of interaction between SpaO and SpaO_C_, we purified constructs covering different regions of SpaO and combined them for native MS and SEC-MALS/SAXS analysis (Table 1). By themselves, both the SpaO N-terminal domain (SpaO_1-145_) and a construct containing the SPOA1 and SPOA2 domains (SpaO_140-297_) are mostly monomeric in solution (Fig. S4A, Table S4, SASDC88 and SASDEK7). Combination of these proteins with SpaO_C_ in both SEC-MALS/SAXS and native MS subsequently showed the formation of a stable SpaO_1-145_-2SpaO_C_ complex resembling the SpaO:2SpaO_C_ stoichiometry, while only very low levels of complexes between SpaO_C_ and SpaO_140-297_ could be detected (Fig. 3A-C, Fig. S4C, D, Table S4 and SASDC98). We characterized the interaction between SpaO_1-145_ and the SpaO_C_ dimer by isothermal titration calorimetry (ITC) and obtained a K_d_ of 1.04 ± 0.21 μM, which also demonstrates strong affinity between these proteins (Fig. 3D, E). Together, these results demonstrate that the intermolecular interaction between the SpaO isoforms is mediated by SpaO_C_ stably binding to the N-terminal domain of SpaO. Furthermore, no significant interaction was detected between the N-terminal domain and the C-terminal SPOA1-SPOA2 domain dimer of SpaO (Fig. 3C, Fig. S4B), indicating that in SpaO these domains are held together largely by their covalent linkage, suggesting conformational flexibility between them.

**Figure 3.**
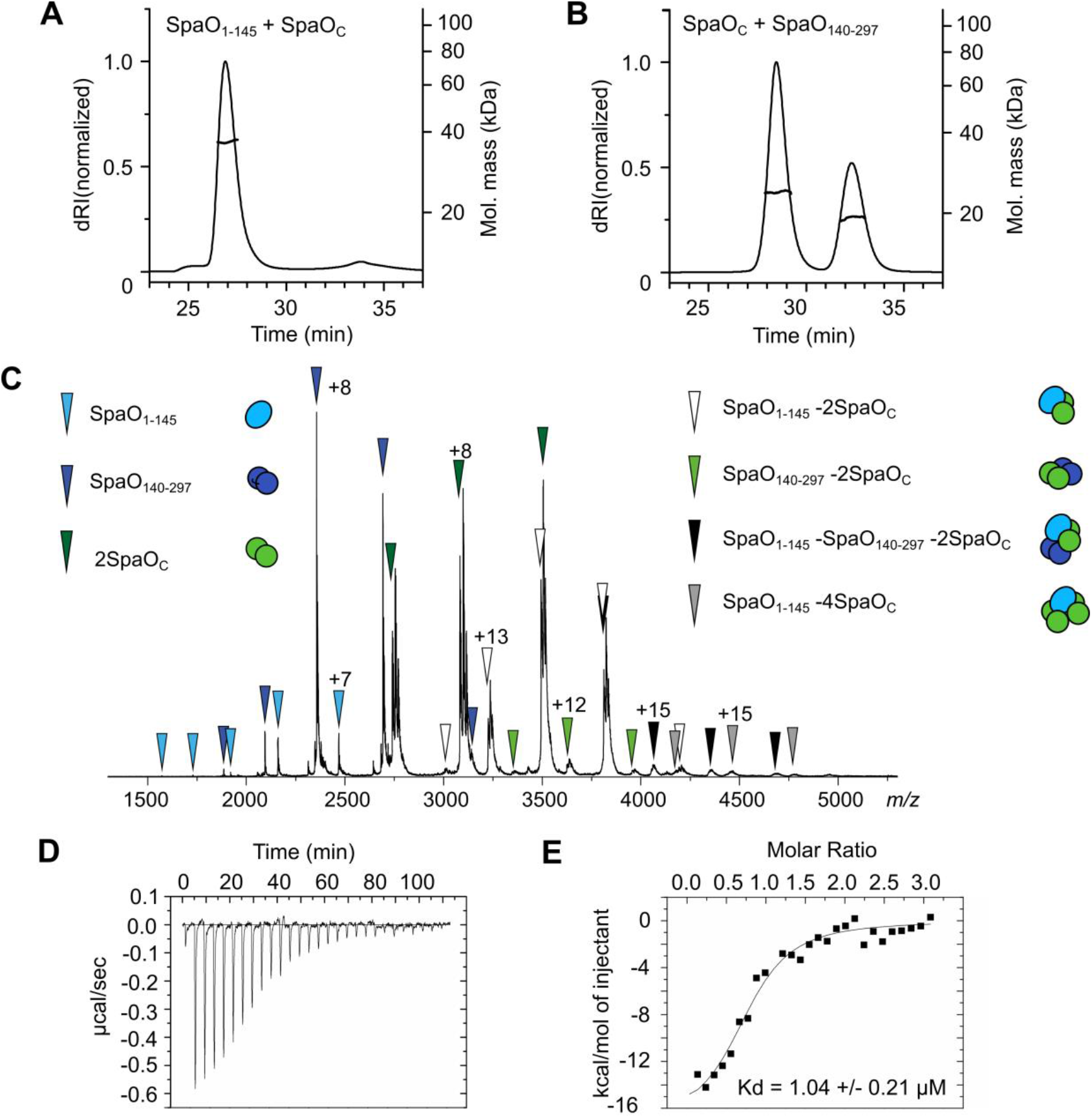
Analysis of inter- and intramolecular domain interactions in SpaO-2SpaO_C_. (A) SEC-MALS analysis of co-purified SpaO_1-145_/SpaO_C_. SEC elution profiles (dRI traces) and the weight-averaged molar masses across the elution peaks are shown. The experimental mass is consistent with the formation of SpaO_1-145_-2SpaO_C_ complexes (theoretical mass of 42kDa). (B) SEC-MALS analysis of combined SpaO_140-297_ and SpaO_C_. (C) Analysis of interactions between SpaO domains by native MS of mixed SpaO_1-145_, SpaO_140-297_ and SpaO_C_. Besides monomeric components, SpaO_1-145_-2SpaO_C_ heterotrimers (white arrows) were found. Other species like SpaO_1-145_-2SpaO_C_-SpaO_140-297_ heterotetramers (black arrows), SpaO_1-145_-4SpaO_C_ (grey arrows) and 2SpaO_C_-SpaO_140-297_ (light green arrows) were detected at very low levels. The used SpaO_C_ sample comprised two protein species with a difference of about 131 Da, resulting in a characteristic peak fine structure with three distinct maxima for complex species containing 2 SpaO_C_. Native mass spectra of SpaO_C_ mixed with only SpaO_1-145_ or SpaO_140-297_ can be found in Fig. S4C and D. Experimental and theoretical molecular masses are given in Table S3. (D) Analysis of SpaO_C_ and SpaO_1-145_ interaction by isothermal titration calorimetry (ITC). Raw heat signal for 10 μl injections of SpaO_C_ dimer (120 μM) to 1.4 ml of SpaO_1-145_ (8 μM). (E) ITC integrated heats and fits to a 1:1 binding model where SpaO_C_ is considered a dimer. Data shown is representative of two experiments.

**Table 1.** Summary of interacting proteins and domains and complex stoichiometries.

| Interactions | Pull down | Native MS | SEC-MALS | ITC |
| --- | --- | --- | --- | --- |
| SpaO/SpaOc | + | 1:2 (2:4) | 1:2 (2:4) | ND |
| SpaO<sub>V203A</sub>/SpaOc | + | ND | ND | ND |
| SpaO<sub>C</sub>/SpaOc | ND | 1:1 (2:2, 4:4) | 1:1 | ND |
| SpaO<sub>1-145</sub>/SpaOc | + | 1:2 (1:4) | 1:2 | K<sub>d</sub> = 1.04 ± 0.21 μM |
| SpaO<sub>1-145</sub>/SpaO<sub>140-297</sub> | ND | - | - | ND |
| SpaO<sub>140-297</sub>/SpaOc | ND | (1:2) | - | ND |
| SpaO/SpaOc/OrgB | + | 2:4:2 (1:2:2, 1:2:1) | ND | ND |
| SpaO/SpaOc/OrgB<sub>1-105</sub> | + | ND | ND | ND |
| SpaO/SpaOc/OrgB<sub>106-226</sub> | +/- | ND | ND | ND |
| SpaO<sub>V203A</sub>/OrgB | + | Not stable | ND | ND |
| OrgB/InvC | + | Not stable | ND | ND |
| OrgB<sub>1-105</sub>/InvC | +/- | ND | ND | ND |
| OrgB<sub>106-226</sub>/InvC | + | ND | ND | ND |
| OrgB/InvC<sub>1-79</sub> | + | ND | ND | ND |
| OrgB/InvC<sub>80-431</sub> | - | ND | ND | ND |
| SpaO/SpaOc/OrgB/InvC | + | 1:2:2:1, 2:4:2:1 (2:2:2:1) | 2:4:2:1 | ND |

Alternative complexes detected by native mass spectrometry (MS) and SEC-MALS are indicated in brackets. ND= Not determined.

### Sorting platform subcomplexes of SpaO_C_, SpaO, OrgB and InvC are stable in solution

Next, we determined the interactions of SpaO and SpaO_C_ with other proteins of the *Salmonella* sorting platform in solution by co-expression with the interaction partners OrgB and InvC (8) in *E. coli* (Table 1). While OrgB by itself was insoluble (data not shown), stable complexes of SpaO/SpaO_C_/OrgB, SpaO/SpaO_C_/OrgB/InvC and OrgB/InvC were soluble and could be purified for further characterization (Fig. 4A). We also tested co-expression of these proteins with OrgA, but this did not yield any soluble OrgA-containing complexes, suggesting that OrgA could either require the presence of needle base-forming proteins for correct folding, or stay localized to the membrane or needle base and not participate in soluble sorting platform complexes.

**Figure 4.**
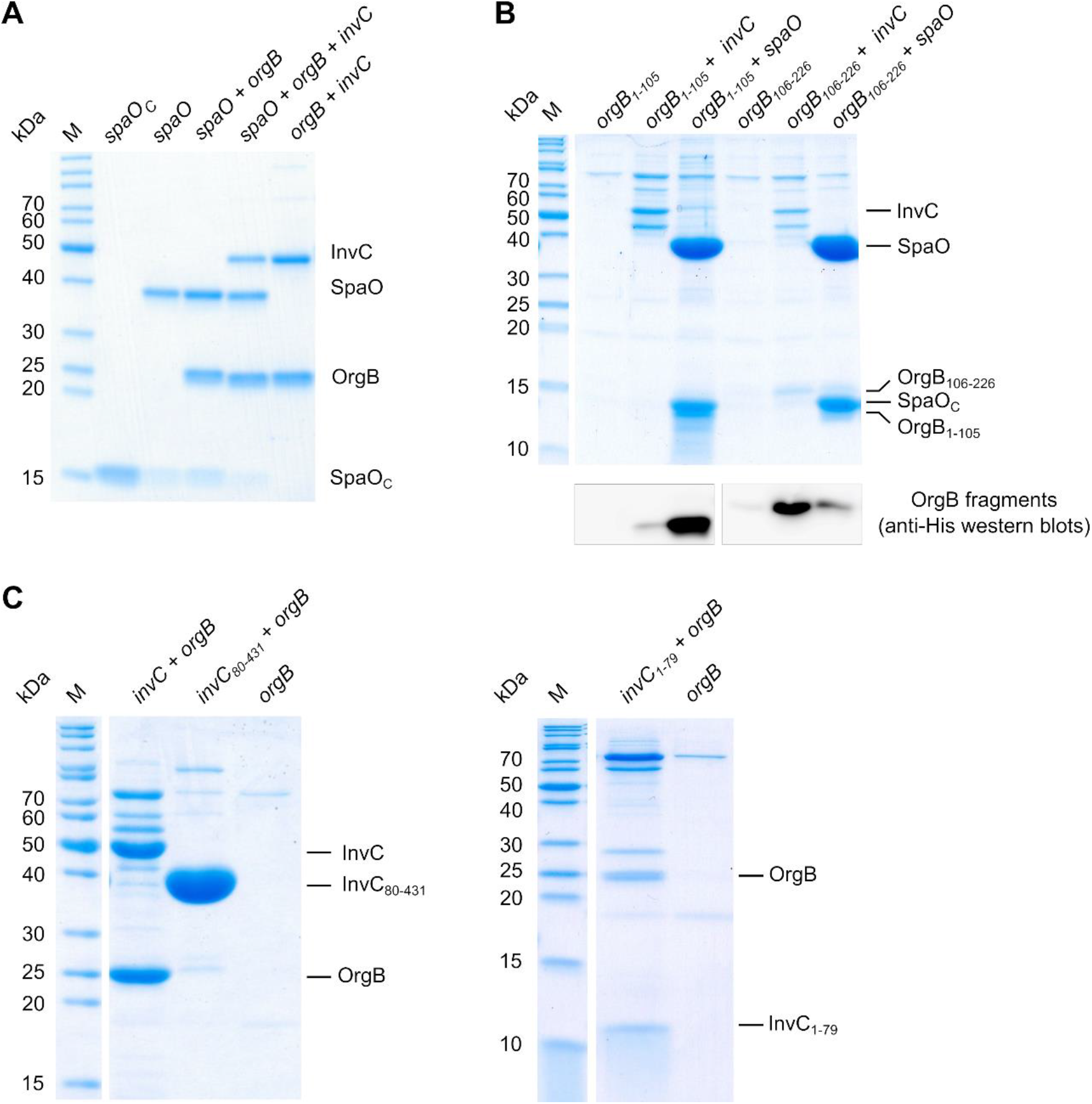
OrgB interacts with SpaO and InvC to form stable sorting platform subcomplexes (A) Coomassie-stained SDS-PAGE of sorting platform proteins co-expressed in *E. coli* and purified by affinity purification and size-exclusion chromatography. Co-expressed genes are indicated above the gel, molecular mass markers are included in lane M. Affinity purification was achieved by use of C-terminal *Strep-*tags for *SpaO_C_* and *spaO*, and a C-terminal *Strep-*tag on InvC for *spaO+orgB+invC* and *orgB+invC*. In the case of *spaO+orgB* affinity purification involved two steps using both a C-terminal His-tag on OrgB and a C-terminal *Strep*-tag on SpaO/SpaO_C_. (B) Top: Coomassie-stained SDS-PAGE of OrgB fragments co-expressed with *Strep*-tagged *invC* or *spaO* in *E. coli* and purified by *Strep*-Tactin affinity purification. Bottom: detection of the His-tagged OrgB fragments by western blot. (C) Coomassie-stained SDS-PAGEs of *Strep*-tagged InvC fragments co-expressed with OrgB and purified by *Strep*-Tactin affinity purification.

In order to determine the regions of OrgB involved in interactions with SpaO/SpaO_C_ and InvC, we dissected OrgB into its N-terminal (residues 1-105) and C-terminal (residues 106-226) halves and co-expressed His-tagged variants of these together with *Strep*-tagged InvC or SpaO/SpaO_C_. Subsequent *Strep*-Tactin affinity purification showed that OrgB_1-105_ is pulled down by SpaO/SpaO_C_, while only trace amounts co-purified with InvC (Fig. 4B). Conversely, OrgB_106-226_, was pulled down by InvC, and even though small amounts could also be pulled down by SpaO/SpaO_C_, the ratio between SpaO/SpaO_C_ and OrgB_106-226_ indicates that the affinity between them is low. We similarly dissected InvC after residue 79 and tested the ability of *Strep*-tagged InvC1-79 and InvC_80-431_ to pull down OrgB. While both full-length InvC and InvC1-79 were able to co-purify OrgB, this was not the case for InvC_80-431_ (Fig. 4C), showing that the N-terminal 79 amino acids of InvC are both necessary and sufficient for its interaction with OrgB. Together, these results show that OrgB interacts through its C-terminus with the N-terminal 79 amino acids of InvC and confirm that the binding site for SpaO is located in the N-terminus of OrgB (15).

### OrgB dimers induce dimerization of SpaO-2SpaO_C_

We analyzed the SpaO/SpaO_C_/OrgB complex by native MS and found that the major molecular species contains two units of SpaO-2SpaO_C_ bound to two molecules of OrgB, resulting in 2(SpaO-2SpaO_C_)-2OrgB complexes. Less abundant species, possibly representing assembly intermediates of this complex, were also identified (Fig. 5). The vast majority of OrgB-containing complexes possesses two molecules of OrgB, which indicates that OrgB exists mainly in dimeric form, similar to its flagellar homolog FliH (25). When we subjected the 2(SpaO-2SpaO_C_)-2OrgB species to MS/MS analysis, a single OrgB dissociated from the complex, while a second, less prominent dissociation pathway led to the dissociation of a SpaO monomer (Fig. S5A).

**Figure 5.**
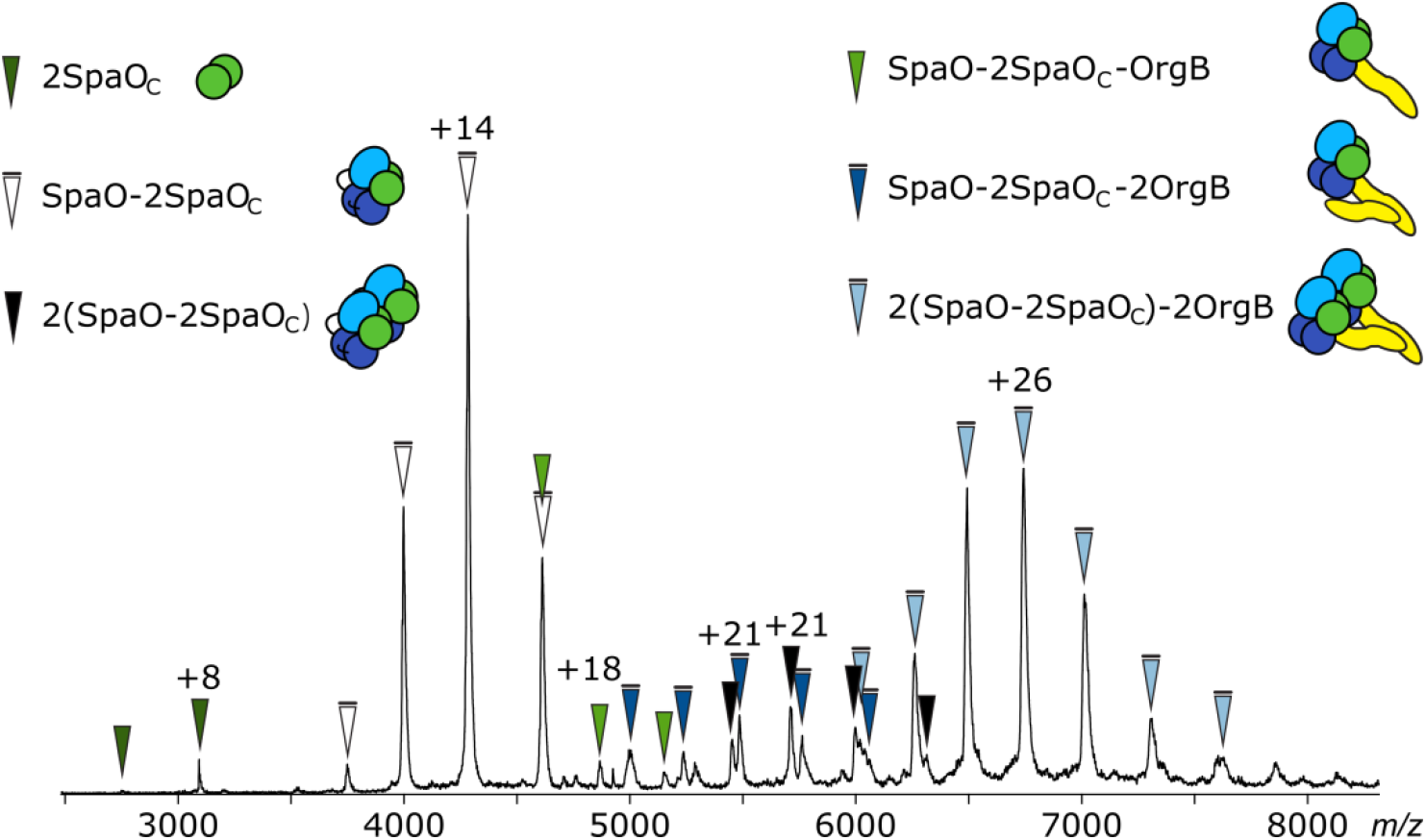
Native mass spectrum of SpaO/SpaO_C_/OrgB complexes. SpaO-2SpaO_C_ heterotrimers (white arrows) bind to OrgB dimers resulting in 2(SpaO-2SpaO_C_)-2OrgB complexes (light blue arrows). 2(SpaO-2SpaO_C_) heterohexamers (black arrows) and a small fraction of SpaO-2SpaO_C_ complexes bound to OrgB monomers (light green arrows) and dimers (dark blue arrows) were also observed. Experimental and theoretical molecular masses are given in Table S3.

We also attempted native MS analysis of the mutant SpaO_V203A_, in which the start codon of SpaO_C_ has been mutated, both alone and in complex with OrgB. While it was possible to purify both SpaO_V203A_ and SpaO_V203A_/OrgB complexes lacking SpaO_C_ by affinity purification and SEC, these complexes were unstable and could not be detected in native MS or successfully analyzed by other methods. This highlights the importance of SpaO_C_ not only for the stability of SpaO, but also of higher-order sorting platform complexes containing OrgB.

### The ATPase InvC binds to SpaO/SpaO_C_/OrgB complexes to form the core building block of the sorting platform

The most comprehensive sorting platform subcomplexes we obtained in this study contained the four proteins SpaO, SpaO_C_, OrgB and InvC. Native MS revealed that the ATPase InvC is present in different types of complexes, with species of SpaO-2SpaO_C_-2OrgB-InvC and 2(SpaO-2SpaO_C_)-2OrgB-InvC stoichiometry being the most abundant in the spectra (Fig. 6A). Complexes of 2OrgB-InvC and 2SpaO-2SpaO_C_-2OrgB-InvC stoichiometry were detected at lower levels. Importantly, InvC was detected exclusively in complexes containing OrgB dimers, which is in agreement with our findings that the OrgB C-terminal region binds to the N-terminus of InvC (Fig. 4B, C) and previously reported CET maps and pull-down assays (10,15). It should be noted that the signal intensity ratios of the different complex species were heavily dependent on the electrospray conditions and while the presented spectrum was selected for a high resolution, the majority of acquired spectra showed higher signal intensities for high molecular weight complexes. However, direct translation of signal intensity ratios into complex ratios in solution is not possible due to fluctuating signals and different ionization and transmission efficiencies of different complex species. Nevertheless, the different observed species indicate a degree of dynamic association and dissociation of subunits within the system. In addition to the species identified in the presented mass spectrum, occasionally signals in the higher *m/z*-range were observed, depending on the electrospray conditions (Fig. S6). Due to the low resolution and signal intensity, charge states for these peaks could not be unambiguously identified in MS or MS/MS measurements. However, the mass range and peak interval suggest the presence of complexes with masses of approximately 433 kDa, possibly dimers of 2(SpaO-2SpaO_C_)-2OrgB-InvC.

**Figure 6.**
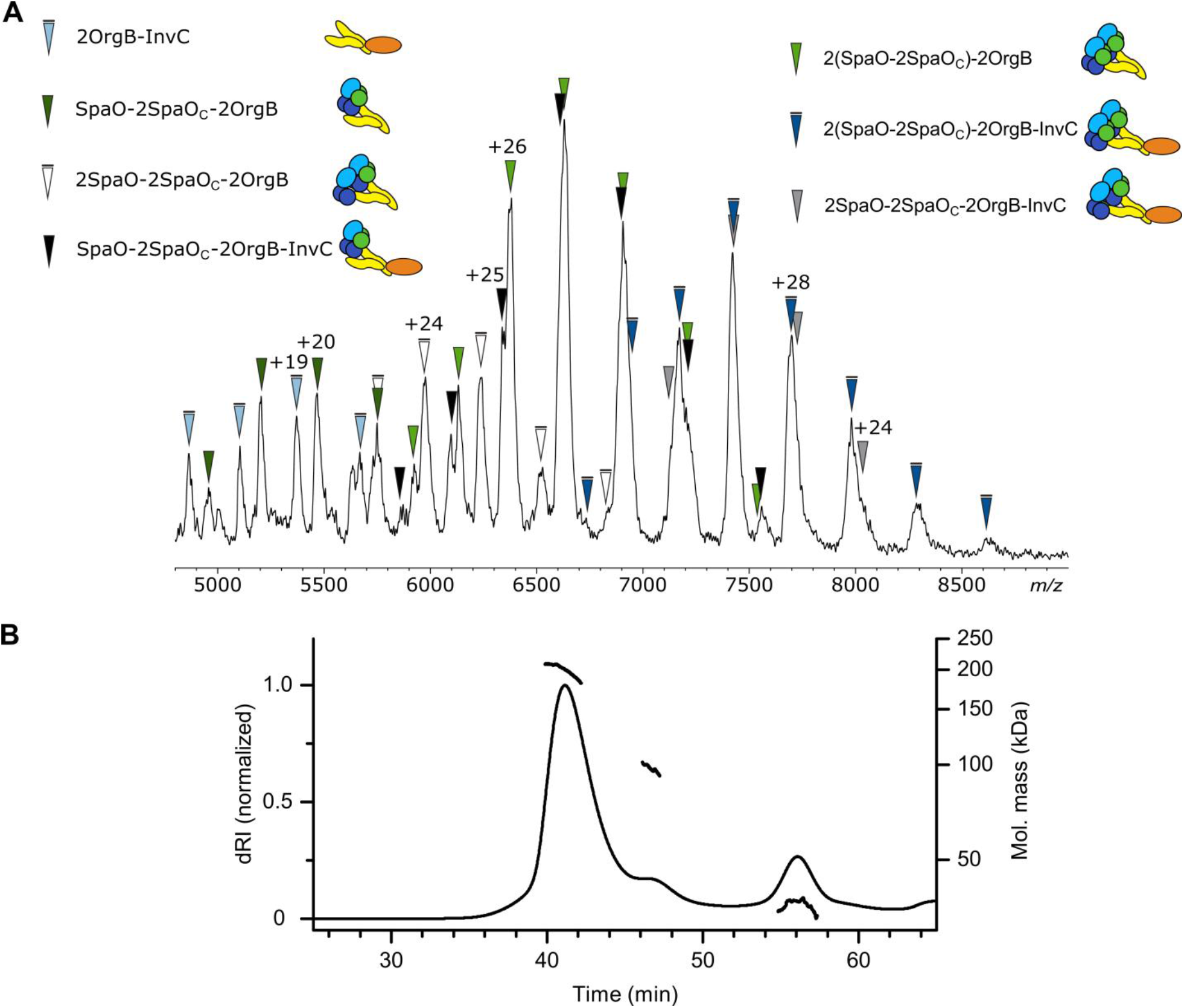
The ATPase InvC binds to SpaO/SpaO_C_/OrgB complexes. (A) Native mass spectrum of SpaO/SpaO_C_/OrgB/InvC complexes. InvC-containing complexes of 2OrgB-InvC (light blue arrows) SpaO-2SpaO_C_-2OrgB-InvC (black arrows), 2(SpaO-2SpaO_C_)-2OrgB-InvC (dark blue arrows) and 2SpaO-2SpaO_C_-2OrgB-InvC (grey arrows) are observed. Experimental and theoretical molecular masses are given in Table S3. (B) SEC-MALS analysis of SpaO/SpaO_C_/OrgB/InvC complexes. SEC elution profiles (dRI traces) and the weight-averaged molar masses across the elution peaks are shown.

In CID MS/MS measurements of the different identified SpaO/SpaO_C_/OrgB/InvC complexes the dissociation of a single OrgB monomer was observed in every case (Fig. S5B, C). Because no other components were lost together with the OrgB, this dissociation pattern allows us to conclude that the interactions of the OrgB dimer with both SpaO/SpaO_C_ and InvC are mediated by the same OrgB molecule, while the other is less tightly integrated in the complex.

We further characterized the SpaO/SpaO_C_/OrgB/InvC complexes using SEC-MALS and SEC-SAXS. MALS revealed a molecular mass range over the main SEC elution peak of approximately 208 to 180 kDa (Fig. 6B), which is in good agreement with the complexes identified in native MS (Table S3). Since this analysis showed the later regions of the elution peak to be a mixture of several molecular species, we only used the largely homogenous first half of the peak in the SAXS analysis in order to generate a model with minimal averaging between different species. Subsequently, bead model reconstruction from the SAXS data showed that the SpaO/SpaO_C_/OrgB/InvC adopts an extended L-shape in solution. (Fig. 7A-C, Table S4, SASDEJ7). By simultaneously employing the SAXS data of the SpaO/SpaO_C_/OrgB/InvC and the SpaO-2SpaO_C_ complexes in a multiphase bead modeling approach (26), the position of SpaO-2SpaO_C_ within the larger complex could be determined. The resulting multiphase bead model indicates that SpaO-2SpaO_C_ is located in the shorter leg of the extended L-shape (Fig. 7D).

**Figure 7.**
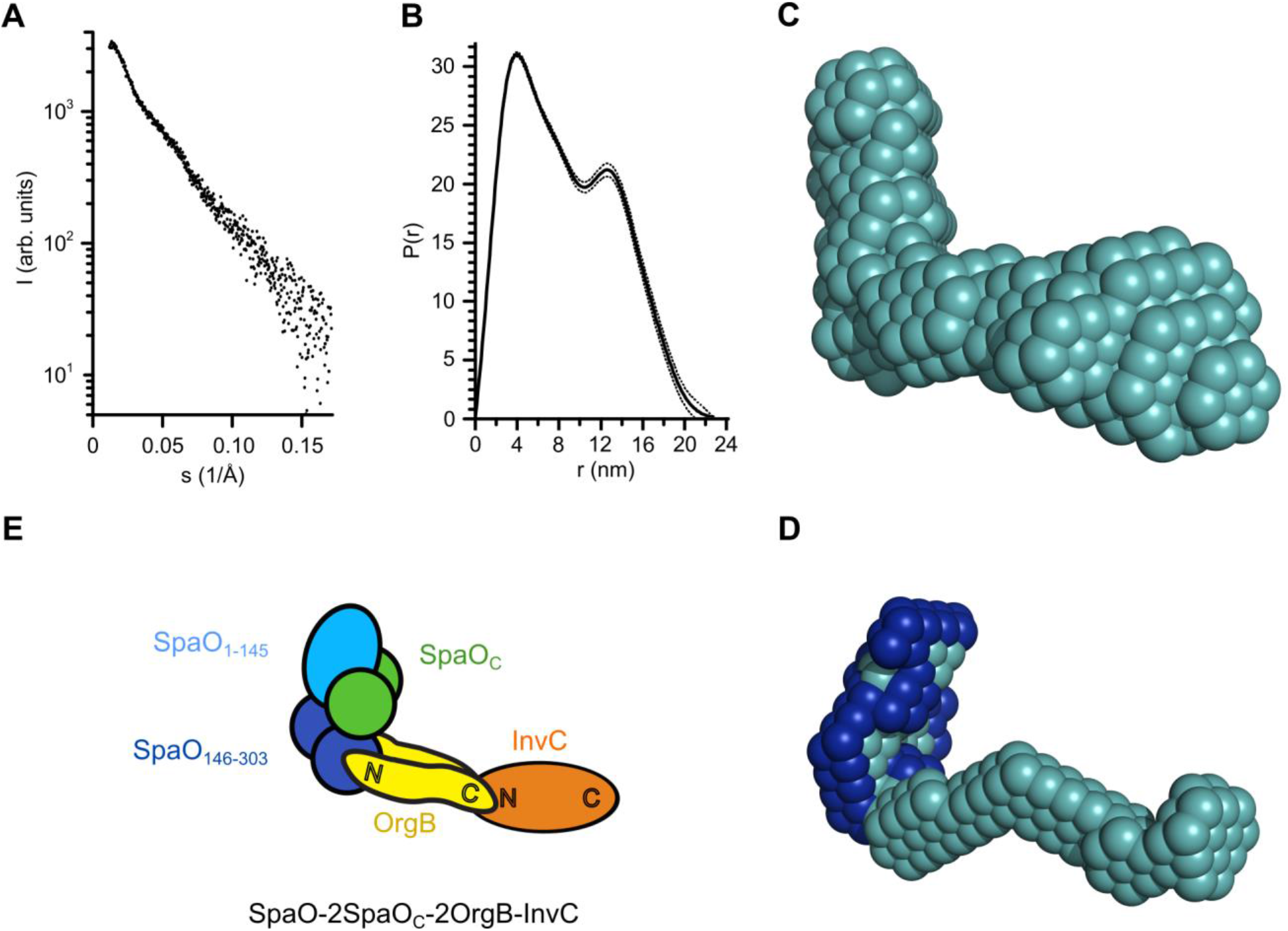
Analysis of SpaO/SpaO_C_/OrgB/InvC complexes by small-angle X-ray scattering. (A) Small-angle X-ray scattering profile of SpaO/SpaO_C_/OrgB/InvC. (B) Pair-distance distribution function *P*(*r*) computed from the SAXS data (A). (C) SAXS-based *ab initio* bead model of SpaO/SpaO_C_/OrgB/InvC. D) MONSA multi-phase modeling using SAXS data of both of SpaO/SpaO_C_/OrgB/InvC and SpaO/SpaO_C_. The phase corresponding to SpaO/SpaO_C_ is colored dark blue. E) Schematic model of the SpaO/SpaO_C_/OrgB/InvC complex taking into account the association of SpaO_C_ with the SpaO N-terminal domain SpaO_1-145_ (Fig. 3), the interaction between the SpaO SPOA1-SPOA2 dimer and the N-terminus of OrgB (15), and the interaction between the C-terminal domain of OrgB and the N-terminal domain of InvC (Fig. 4B, C).

Unfortunately, due to the complexity of the studied system the generation of a reliable SAXS-based atomistic hybrid model of the SpaO/SpaO_C_/OrgB/InvC complex is hindered by a number of uncertainties, which include the number of different subunits, flexibility of the complex in solution (indicated by Kratky analysis, see SASBDB), the remaining possibility of heterogeneity in the SEC peak region used for SAXS analysis and a lack of high-resolution structure for many of the complex components. Nevertheless, by combining the SpaO/SpaO_C_/OrgB/InvC SAXS data with our native MS results and the interactions between different subunit domains (Table 1), it is possible to construct a schematic model of the architecture of the soluble SpaO/SpaO_C_/OrgB/InvC complex (Fig. 7E). Thus, while SpaO-2SpaO_C_ occupies the shorter leg of the L-shape, InvC-OrgB would be placed in the longer leg with OrgB forming a linker between SpaO-2SpaO_C_ and InvC. Interestingly, even though native MS and MALS indicate the presence of two SpaO-2SpaO_C_ heterotrimers in the SpaO/SpaO_C_/OrgB/InvC complex (Fig. 6, Table S3), both the multiphase analysis (Fig. 7D) and comparison of the SpaO/SpaO_C_ and SpaO/SpaO_C_/OrgB/InvC SAXS bead structures (see SASBDB for details) indicate that only a single SpaO-2SpaO_C_ heterotrimer can be accommodated in the short leg of the L-shape, suggesting that the SAXS structure is that of a complex with SpaO-2SpaO_C_-2OrgB-InvC stoichiometry. This apparent discrepancy could be due to uncertainties in the SAXS bead model caused by flexibility of the complex or heterogeneity in the SEC peak region used for SAXS analysis.

Because the extended SAXS shape of the SpaO-2SpaO_C_-2OrgB-InvC complex is reminiscent of the pod densities seen in the *in-situ* 3D CET map of the *Salmonella* needle complex (10), we hypothesized that this complex represents the soluble core building block from which the full sorting platform is assembled. To test this hypothesis, we superimposed the *ab initio* SAXS bead model with the CET map (Fig. 8), which shows a good correspondence between the two structures and orients the SAXS shape in a way that places SpaO-2SpaO_C_ in the outer pods, InvC in the central hub and 2OrgB in the linker region between the two. This is in good agreement with the assignments by CET using fluorescent protein tags and sorting platform protein deletions (10,27). If six units of the SAXS bead model were to be placed within the CET map, steric clashes would occur in the central hub region. Interestingly, Kratky analysis of the SAXS data showed conformational flexibility of the building block complexes, indicating an ability to undergo conformational changes upon assembly of the complete sorting platform. In fact, by rotating InvC in our model upwards by 90° around its interaction site with OrgB and shifting that interaction site towards the bottom of the central hub, InvC would be re-oriented into a configuration parallel to the outer pods with its C-terminus pointing towards the T3SS basal body. This change would both resolve the steric clashes and allow for the formation of an InvC ATPase hexamer to fill the central hub region of the CET map.

**Figure 8.**
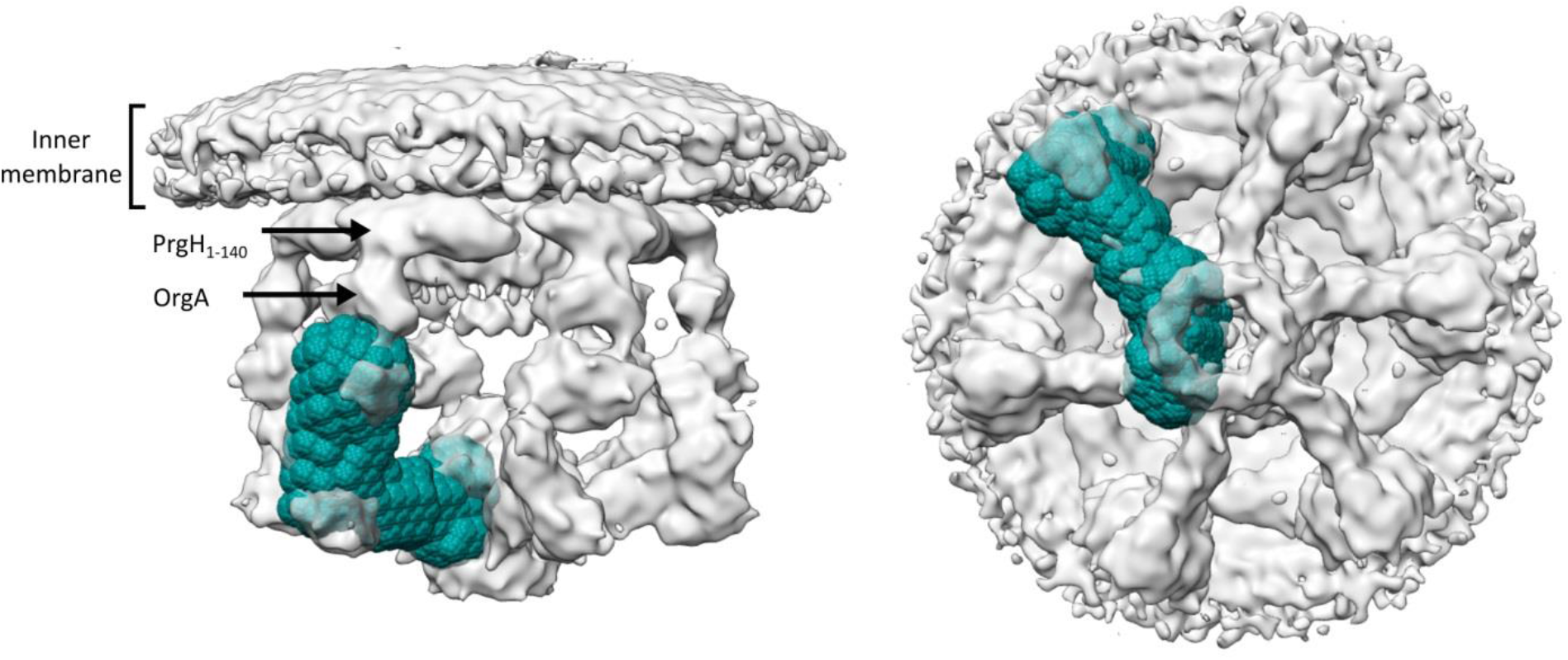
Superposition of the SAXS-based bead model (cyan) with the *in-situ* CET structure of the *Salmonella* Typhimurium sorting platform (EMDB ID: EMD-8544, grey). Shown are a side view (left) and bottom view (right). The N-terminal domain of PrgH (PrgH_1-140_) and OrgA are labeled according to Hu et al., 2017 (10).

It should, however, be noted that this *in silico* approach relies on the superposition of two structures of low resolution, both of which are associated with their own errors, posing a limit on the conclusions that can be drawn from it. Therefore, while the good agreement between our SAXS structure and the CET map supports our idea that the SpaO/SpaO_C_/OrgB/InvC complex represents the soluble core building block of the sorting platform, biochemical studies will be required to show the assembly of these soluble complexes into the complete sorting platform.

## Discussion

The sorting platform, together with the export apparatus complex, is still one of the less well characterized components of the T3SS. In this work we present an analysis of inter-subunit interactions, stoichiometry and shape of the main soluble module of the SPI-1 sorting platform in *S.* Typhimurium. Expression and functional analysis confirm that the gene encoding the protein SpaO produces an additional short protein SpaO_C_ that comprises the C-terminus of the SpaO sequence, and that spaO_FL_ is essential for type III secretion, while SpaO_C_ appears non-essential but required for full secretion efficiency (16,17). Interestingly, a similar phenotype has been observed for the *Salmonella* SPI-2 orthologue protein SsaQC (18) and the remaining secretion activity upon deletion of the shorter protein product appears to be unique to the two T3SSs of *Salmonella.* This raises the possibility that cross-complementation might occur between the *Salmonella* T3SSs, a hypothesis that could be the subject of future investigations. On the other hand, given that MALDI MS showed that low levels of a SpaO_C_-like protein were still produced from a *spaO* variant carrying a mutation in the SpaO_C_ start codon (*SpaO_V203A_*), it is possible that the incomplete loss of secretion and invasion activity of the *ΔSpaO_C_* mutant might be due to partial complementation by such a product that appears to be produced by proteolysis even in the absence of internal translation initiation (Fig. 1B, C). Overall, these data are consistent with the results of previous studies in the *Salmonella* SPI-1 and SPI-2 systems, as well as *Shigella* and *Yersinia* (12,16–20), suggesting that the alternative translation into a full-length protein and a shorter product may represent a widespread strategy among the SctQ proteins of virulence-associated T3SSs.

The isoform SpaO_C_ forms a homodimer that binds to the full-length SpaO to form SpaO-2SpaO_C_ complexes, similar to the 1:2 complexes observed for the *Shigella* Spa33 and the *Yersinia* YscQ homologs (19,20). However, while the Spa33-2Spa33_C_ trimers readily assembled into higher-order oligomers, we found only little dimerization of trimers and no further oligomerization for SpaO-2SpaO_C_. Additionally, our analysis shows that the SpaO_C_ dimer stably associates with the N-terminal domain of SpaO (SpaO_1-145_). In contrast, stable interactions between SpaO_C_ and the SPOA domains of SpaO, like those observed for the homolog Spa33, could not be detected (20). Interestingly, in a recent study a SpaO variant carrying a photo-activatable amino acid in the SPOA2 domain (residue 289) was found to cross-link with SpaO_C_, indicating interaction between these regions after all (16). However, given the irreversibility of cross-linking and the comparatively low levels of cross-linked species in that study, these might have been the result of more transient interactions such as those indicated by the low levels of SpaO_140-297_-SpaO_C_ complexes observed in native MS (Fig. 3C). Importantly, our newly found stable interaction between the N-terminal domain of SpaO and SpaO_C_ has implications for any model of the structure of the T3SS cytosolic complex, which is currently based on interactions between the small SctQ protein isoform and the SPOA1-SPOA2 domain dimer of the full-length variant.

The SpaO-2SpaO_C_ heterotrimer interacts with the ATPase regulator OrgB to form stable 2(SpaO-2SpaO_C_)-2OrgB complexes (Fig.5) and we therefore propose that OrgB exists as a dimer comparable to its flagellar homolog FliH (25). Interestingly, while it has been suggested that the binding of SpaO_C_ and OrgB to SpaO may be mutually exclusive due to overlapping binding sites on the C-terminal SPOA1-SPOA2 domains of SpaO (15,16), our data shows that SpaO can simultaneously interact with both of these proteins. This is consistent with our finding that SpaO_C_ interacts with the N-terminal domain of SpaO rather than the C-terminal SPOA domains. In addition, in CID MS/MS of 2(SpaO-2SpaO_C_)-2OrgB complexes the dissociation of both SpaO and OrgB monomers was observed, indicating that the recruitment of OrgB leads to a stabilization of SpaO_C_ within the complex. While our data does not offer a clear mechanism for this stabilization, it is conceivable that it involves direct interactions between SpaO_C_ and OrgB that occur in addition to those of the extreme N-terminus of OrgB and the SpaO SPOA1-SPOA2 dimer (15). Furthermore, the dissociation of either OrgB or SpaO without the simultaneous loss of other subunits suggests that direct interactions between the two SpaO-2SpaO_C_ trimers are promoted in these complexes. It should be noted that the observed MS/MS dissociation pattern is also compatible with a complex architecture in which both SpaO-2SpaO_C_ heterotrimers are associated with the same OrgB unit. However, this arrangement seems unlikely since an association of one SpaO-2SpaO_C_ trimer to one OrgB would be expected in light of the reported interaction between the N-terminus of OrgB and the SPOA1-SPOA2 domains of SpaO (15). Nevertheless, it cannot be excluded given the asymmetry of OrgB units within the OrgB dimer revealed by MS/MS of SpaO/SpaO_C_/OrgB/InvC complexes (see below).

Complexes of SpaO, SpaO_C_ and OrgB associate with the ATPase InvC to form both SpaO-2SpaO_C_-2OrgB-InvC and 2(SpaO-2SpaO_C_)-2OrgB-InvC complexes, in which OrgB acts as a central connector by binding of its N-terminus to SpaO-2SpaO_C_ and its C-terminus to InvC. Based on MS/MS experiments (Fig. S5), we propose that both of these interactions are formed by the same OrgB subunit, while the second OrgB is less tightly integrated in the complex, possibly acting to stabilize extended helical regions in the first OrgB. SAXS analysis showed that the SpaO/SpaO_C_/OrgB/InvC complexes adopt an extended L-shaped structure in solution. Because this conformation is in good agreement with the *in situ* cryo-electron tomography (CET) structure of the *Salmonella* SPI-1 T3SS (10), we propose that the SpaO/SpaO_C_/OrgB/InvC complexes identified in this study represent the main soluble building blocks of the sorting platform. These complexes would bind to other T3SS proteins like the docking protein OrgA, InvI or the export apparatus and undergo a conformational change in re-orienting the ATPase InvC to assemble the complete sorting platform at the base of the T3SS needle complex. Adding to the similarity of the SAXS and CET structures the fact that we found SpaO_C_ in all of the sorting platform subcomplexes, its importance for their stability and that SpaO_C_ is itself stabilized in these complexes by the presence of OrgB, it can be hypothesized that SpaO_C_ is an integral structural part of the sorting platform similar to the *Yersinia* homolog YscQC, which has previously been shown to co-localize with YscQFL into sorting platform complexes at the bacterial membrane (12). On the other hand, the *in vitro* nature of our study means that it cannot be excluded that SpaO_C_ plays a role in the soluble forms of the building blocks and might dissociate from the complexes upon assembly of the complete sorting platform *in vivo*, as has been suggested by the lack of additional densities in CET maps of sorting platforms from strains expressing SpaO_C_ fused to a fluorescent protein (16).

The superposition between our SAXS bead model and the CET map suggests that the individual legs as seen by tomography would be of SpaO-2SpaO_C_-2OrgB-InvC stoichiometry, bringing the assembled sorting platform to 6SpaO-12SpaO_C_-12OrgB-6InvC. While these numbers are compatible with the stoichiometry determined by fluorescence microscopy for InvC and OrgB, SpaO has been indicated to be present in the sorting platform at a higher copy number of approximately 24 (13). Our findings show that the soluble building blocks can recruit an additional SpaO-2SpaO_C_ trimer and it is conceivable that further units dynamically associate with the sorting platform at the T3SS needle base. In fact, the dynamic exchange of the SpaO homolog YscQ in *Yersinia* has previously been observed by fluorescence microscopy and found to increase during the active secretion process (12). Together with the observation that the diffusion behavior of cytosolic populations of sorting platform components also changes upon secretion activation, this indicates that soluble sorting platform complexes might play an important role in the function of type 3 secretion (28). Thus, it can be hypothesized that soluble building blocks of SpaO/SpaO_C_/OrgB/InvC could act as T3SS substrate shuttles that recruit substrate-chaperone complexes in the cytosol and transfer them to the basal-body associated sorting platform for subsequent secretion. Furthermore it can be speculated that the dissociation of SpaO/SpaO_C_/OrgB from hexameric T3SS-associated InvC might act to fully activate the secretion process by overcoming the inhibitory effect of OrgB on InvC ATPase activity (25,29).

## Experimental Procedures

### Cloning and mutagenesis of *Salmonella* genes

Genes ligated into the expression vectors pASK-IBA (IBA GmbH, Göttingen, Germany), pET (Novagen, Madison, WI, USA), or pCDFDuet-1 (Novagen, Madison, WI, USA) were derived from *Salmonella* Typhimurium strain SL1344 using standard techniques. All PCRs were performed using Phusion polymerase (New England Biolabs, Ipswich, MA, USA) and oligonucleotides synthesized by Sigma-Aldrich or Eurofins Genomics. Site-directed mutagenesis of the *spaO* gene was performed according to the QuikChange PCR site-directed mutagenesis protocol (Agilent, Santa Clara, CA, USA). All primers used in this study can be found in Table S5.

*Salmonella* genomic *spaO* deletion was carried out by homologous recombination using the λ Red recombinase system (30). Briefly, the λ Red recombinase plasmid pKD46 was expressed in *S.* Typhimurium SL1344 and a kanamycin cassette flanked by two 50bp regions homologous to the *spaO* gene was subsequently transformed into the strain for homologous recombination. The Δ*SpaO_C_*, Δ*spaO*_FL_, *spaO*-3xFLAG, Δ*SpaO_C_*-3xFLAG and Δ*spaO*_FL_-3xFLAG strains were generated following a similar protocol, introducing a tetracycline cassette into the *spaO* region as described above. In a second step, the tetracycline cassette was replaced by *spaO* DNA carrying mutations and colonies were selected on tetracycline-sensitivity selection media (31,32). To generate the Δ*SpaO_C_* strain, silent mutations at the internal putative Shine-Dalgarno region (position 594 to 600, AGGGGGA to gGGcGGc) and start codon (position 607-609, GTG to GTt) of *spaO* were introduced, while the Δ*spaO*_FL_ strain was produced by introducing nonsense mutations shortly after the start codon of *spaO* at amino acid position 28 and 29. For the generation of the strains *spaO*-3xFLAG, Δ*SpaO_C_*-3xFLAG and Δ*spaO*_FL_-3xFLAG, a 3xFLAG-tag was inserted at the C-terminus of *spaO* in the chromosome. Introduction of mutations was verified by PCR and DNA sequencing. The Δ*spi-1* strain was kindly provided by the lab of Arturo Zychlinsky.

### Detection of SpaO and SpaO_C_ in *Salmonella* cells

*spaO*-3xFLAG, Δ*SpaO_C_*-3xFLAG and Δ*spaO*_FL_-3xFLAG strains were grown in LB medium (Luria/Miller) at 37 °C to an OD_600_ of 1. Total cell lysates were separated by SDS-PAGE and analyzed by western blot using anti-FLAG M2 primary antibody (Sigma-Aldrich, St. Louis, MO, USA), horseradish peroxidase (HRP)-conjugated secondary antibodies (Jackson ImmunoResearch Laboratories, West Grove, PA, USA) and ECL western blotting substrates (Thermo Fischer Scientific, Waltham, MA, USA) for protein detection.

### Recombinant gene expression and protein purification

Constructs used for recombinant gene expression in *E. coli* BL21 (DE3) are listed in Table S6. Cells were grown in LB with the appropriate antibiotics at 37 °C. At an OD_600_ of 0.5, the temperature was reduced to 20 °C and gene expression induced by addition of 200 μg/l anhydrotetracycline (AHT, Sigma-Aldrich, St. Louis, MO, USA) for pASK-IBA vectors and/or 0.5 mM IPTG for pET and pCDFDuet plasmids. Cells were grown further for 18 h and harvested by centrifugation.

All purification steps were performed at 4 °C. To purify SpaO_C_, SpaO_1-145_, SpaO_140-297_, SpaO_1-145_/SpaO_C_, SpaO/SpaO_C_, SpaO/SpaO_C_/OrgB/InvC and OrgB/InvC complexes, cell pellets were resuspended in buffer B1 (100 mM Tris pH 7.5, 150 mM NaCl) supplemented with complete EDTA-free protease inhibitor cocktail (Roche), 1 mg/ml lysozyme, 10 μg/ml DNase I and 2 mM 2-mercaptoethanol (2ME). Cell lysis was achieved by French press and lysates were clarified by centrifugation at 48,000 × g for 30 min. The protein complexes were purified by *Strep*-Tactin affinity chromatography and eluted with buffer B1 supplemented with 7.5 mM desthiobiotin. Affinity-purified proteins were polished by size-exclusion chromatography (SEC) on Superdex 75 or Superdex 200 columns (GE Healthcare, Chicago, IL, USA) equilibrated with buffer B2 (20 mM HEPES pH 7.5, 350 mM NaCl). The affinity-purified OrgB/InvC and SpaO/SpaO_C_/OrgB/InvC complexes were further purified by SEC on a Superose 6 column equilibrated with 10 mM Tris-HCl pH 8.0, 50 mM NaCl, with InvC/OrgB having been dialyzed against the same buffer before the SEC. For SpaO/SpaO_C_/OrgB complex purification, cells were resuspended in buffer B3 (20 mM sodium phosphate buffer pH 7.4, 500 mM NaCl) supplemented with 40 mM imidazole, protease inhibitors, 1 mg/ml lysozyme, 10 μg/ml DNase I and 2 mM 2ME. The SpaO/SpaO_C_/OrgB complex was immobilized on HisTrap HP columns (GE Healthcare, Chicago, IL, USA), washed with buffer B3 containing 3 mM ATP and 10 mM MgCl2 and eluted with buffer B3 containing 400 mM imidazole. Eluted proteins were diluted three-fold in buffer B1, purified by *Strep*-Tactin affinity chromatography, followed by SEC in a Superdex 200 column equilibrated with buffer B2.

For solubility analysis of sorting platform proteins (Table S6), cells were lysed by sonication (Sonopuls HD 2070, Bandelin, Berlin, Germany), soluble and insoluble fractions were separated by centrifugation and analyzed by SDS-PAGE and western blot using anti-*Strep* (Qiagen, Hilden, Germany) and anti-His (GE Healthcare, Chicago, IL, USA) primary antibodies, HRP-conjugated secondary antibodies and SuperSignal West Dura Extended Duration Substrate (Thermo Fisher Scientific, Waltham, MA, USA) for detection.

For the purification of OrgB fragments with SpaO/SpaO_C_ and InvC, as well as InvC fragments with OrgB, cells were resuspended in buffer B1 supplemented with complete EDTA-free protease inhibitor cocktail (Roche, Basel, Switzerland), 1 mg/ml lysozyme, 10 μg/ml DNase I, 2 mM 2-ME and 1 mM MgCl2. Cells were lysed by sonication, lysates clarified by centrifugation and protein from the soluble fraction purified by *Strep*-Tactin affinity purification. Eluted proteins were analyzed by SDS-PAGE followed by Coomassie-staining or western blot using an anti-His primary antibody (Thermo Fisher Scientific, Waltham, MA, USA), HRP-conjugated secondary antibody (Jackson ImmunoResearch Laboratories, West Grove, PA, USA) and ClarityMax ECL substrate (Bio-Rad, Hercules, CA, USA).

To test the effect of SpaO_C_ on the solubility of mutant SpaO proteins, plasmid constructs (Table S6) were expressed for 3 h at 37 °C in a *Salmonella spaO*-knockout strain (SL1344Δ*spaO*). Harvested cells were resuspended in phosphate-buffered saline and lysed with BugBuster reagent (Novagen, Madison, WI, USA). Soluble proteins were loaded onto *Strep*-Tactin resin and loaded resin was analyzed by SDS-PAGE and Coomassie staining.

### Protein secretion

Strains were grown in LB at 37 °C for 6 h to induce SPI-1 effector protein secretion. Where appropriate, expression was induced with AHT at an OD_600_ of 0.1. Proteins were precipitated from 12-13 ml of filtered culture supernatants by addition of 15% ice-cold trichloroacetic acid (TCA) and centrifugation at 3.200 × *g* for 90 min. Pellets were washed with ice-cold acetone, air-dried and resuspended in 200 mM Tris-HCl (pH 8.0) containing 200 mM NaCl. Samples were loaded onto SDS-PAGE gels and analyzed by Coomassie staining and western blot. Rabbit anti-SipB, anti-SipC, anti-SipD and anti-SopB polyclonal antibodies were raised and applied for detection of T3SS-dependent substrates in western blot analysis. Anti-FliC (kindly provided by Marc Erhardt’s lab) and anti-DnaK antibodies (Stressgen Biotechnologies, San Diego, CA, USA) were used as loading control and lysis control, respectively. HRP-conjugated secondary antibodies and ECL western blotting substrates were used for protein detection.

### Invasion assay

The murine epithelial cell line MODE-K (33) was cultivated in Dulbecco’s modified Eagle’s medium (DMEM). 5 × 10^5^ cells were seeded in 24-well plates and infected with *Salmonella* strains at a multiplicity of infection (MOI) of 10 for 1 h at 37 °C with 5% CO2 in a humidified tissue culture incubator. After treating the cells with 100 μg/ml gentamycin for 1 h, cells were washed with sterile PBS three times. Infected monolayers were lysed with 1 % Triton-X and colony forming units (CFU) were determined by serial dilution and plating. Relative invasion of each strain was calculated by comparison of the CFUs after invasion with those of the inoculum.

### Isothermal titration calorimetry (ITC)

ITC of SpaO_C_ binding to the SpaO_1-145_ was performed using a MicroCal VP-ITC titration calorimeter (Malvern Panalytical, Almelo, Netherlands) calibrated to 25°C. 1.4 ml of SpaO_1-145_ at 8 μM was placed in the sample cell, and the syringe was loaded with 120 μM of SpaO_C_ dimer. Injections of 10 μl were performed with stirring at 310 rpm and the heat of reaction was recorded. Data were analyzed using Origin (OriginLab, Northampton, MA, USA).

### Native mass spectrometry

Purified protein samples were buffer-exchanged into 50 mM ammonium acetate pH 7.5 (SpaO and SpaO fragments), 300 mM ammonium acetate pH 7.5 (SpaO_C_/SpaO/OrgB) or 50 mM ammonium acetate pH 8 (SpaO/SpaO_C_/OrgB/InvC) using Vivaspin^®^ 500 centrifugal concentrators (Sartorius, Göttingen, Germany). SEC-purified proteins were used for all samples but the SpaO/SpaO_C_/OrgB complex, which was affinity-purified. Samples were loaded into home-made gold-coated glass capillaries (34), which were mounted into the nano electrospray ionization (ESI) source of a QToF 2 mass spectrometer (Waters, Manchester, UK, and MS Vision, Almere, the Netherlands) adapted for high-mass experiments (35) and operated in positive ion mode. Capillary and cone voltages of 1.3 to 1.5 kV and 110 to 150 V were applied, respectively. The source pressure was set in the range of 6 to 10 mbar and argon was used as collision gas at 1.7 to 1.9 × 10^−2^ mbar. Acceleration voltages for collision-induced dissociation (CID) were optimized for resolution and minimal complex dissociation. CID tandem mass spectrometry (MS/MS) experiments on protein complexes were performed to confirm mass assignments and deduce topological information by selecting specific precursor peaks for dissociation and ramping acceleration voltages up to 400 V or until the entire precursor signal disappeared. Cesium iodide spectra (25 mg/ml) were acquired on the same day of each measurement and used to calibrate raw data using MassLynx software (Waters, Manchester, UK). Peak series were assigned with MassLynx and Massign (36). Average measured masses of protein complexes, standard deviations of replicate measurements and average full width at half maximum (FWHM) values as a measure of the mass heterogeneity and resolution are listed in Table S3.

### Small-angle X-ray scattering and multi-angle light scattering

Small-angle X-ray scattering (SAXS) measurements were carried out at the beamline P12 (EMBL/DESY, Hamburg, Germany) (37) at the PETRA III storage ring using a Pilatus 2M detector (Dectris, Baden-Dätwil, Switzerland). The SAXS camera was set to a sample-detector distance of 3.1 m, covering the momentum transfer range 0.008 Å^−1^ < *s* < 0.47 Å^−1^*s* = 4π sin(θ)/λ (where 2 θ is the scattering angle and λ=1.24 Å is the X-ray wavelength). For each SAXS measurement, 75-90 μl of affinity-purified protein sample was loaded onto a Superdex 200 Increase 10/300 GL SEC column (GE Healthcare, Chicago, IL, USA) previously equilibrated with 20 mM HEPES pH 7.5, 150 mM NaCl and eluted at 0.5 ml/min. In the case of the SpaO/SpaO_C_/OrgB/InvC complex, Superose 6 10/300 GL (GE Healthcare, Chicago, IL, USA) equilibrated with 10 mM Tris-HCl pH 8.0, 50 mM NaCl and a flow rate of 0.3 ml/min was used. The sample eluting from the SEC column was split into two fractions using a mobile phase-flow splitter. One fraction was directed to the SAXS flow cell and the other into a triple detector array of UV absorption, multi-angle light scattering (MALS, Wyatt MiniDawn Treos), and RI detectors (Wyatt Optilab T-rEX, both Wyatt, Santa Barbara, CA, USA). Only in the case of SpaO/SpaO_C_/OrgB/InvC, independent experiments were run for SAXS and MALS data acquisition. The molecular masses of the separated sample components eluting from the column were estimated by combining the results from light and X-ray scattering with RI and UV absorption measurements. For each sample the scattering profiles over the elution peak, collected with an exposure time of 1 s each, were separated into sample and buffer regions, appropriately averaged and the signal from the buffer was subtracted using CHROMIXS (38).

### SAXS model-free parameters

The radius of gyration Rg and forward scattering intensity I(0) were determined using Guinier analysis (39) and an indirect Fourier transformation approach by the program GNOM (40), the latter also providing maximum particle dimensions Dmax.

### Structural modelling against SAXS data

*Ab initio* models were reconstructed from the scattering data using bead modelling program DAMMIF and multiphase modelling program MONSA (26,41). Ten independent reconstructions were averaged to generate a representative model with the program DAMAVER (42). The average DAMMIF model was also used to calculate the excluded volume of the particle, V_DAM_, from which an independent MW estimate can be derived (empirically, MM_DAM_ ~ V_DAM_/2). Resolutions of the *ab initio* models were computed using a Fourier Shell Correlation (FSC) based approach (43). Ambiguity associated with spherically averaged single-particle scattering was determined using by AMBIMETER (44).

For the comparison between SAXS data and the electron microscopy density map the program Chimera (45) was used to superimpose a bead model based on the *ab initio* SAXS shape with the *Salmonella* T3SS CET map (EMDB ID: EMD-8544). A contour level of 2.53 was used for the CET.

### Accession Codes

The details of the SAXS analysis and the generated models were deposited at the Small-Angle Scattering Biological Data Bank (SASBDB) under the codes: SASDC68 (SpaO_C_); SASDEK7 (SPAO_140-297_); SASDC88 (SpaO_1-145_); SASDC98 (SpaO_1-145_/SpaO_C_); SASDC78 (SpaO/SpaO_C_); SASDEJ7 (SpaO/SpaO_C_/OrgB/InvC).

## Acknowledgements

The authors gratefully acknowledge P. Jungblut and M. Schmidt for the MALDI-TOF/TOF mass spectrometry analysis, B. Jaschok-Kentner and W. Blankenfeldt for Edman sequencing, C. Jeffries for the MALS analysis, M. Lunelli for CET and SAXS alignment and J. de Diego for her useful comments and critical reading of the manuscript.

This work was funded by the European Research Council under the European Community’s Seventh Framework Programme (FP7/2007–2013). The Heinrich Pette Institute, Leibniz Institute for Experimental Virology is supported by the Free and Hanseatic City of Hamburg and the German Federal Ministry of Health. JH and CU are funded by the Leibniz Association through SAW-2014-HPI-4 grant. AT was supported by the EMBL interdisciplinary Postdoc Programme under Marie Curie COFUND Actions.

## Conflicts of Interest

The authors declare that they have no conflicts of interest with the contents of this article.

## References

1. Hueck, C. J. (1998). Type III protein secretion systems in bacterial pathogens of animals and plants. Microbiol. Mol. Biol. Rev. 62, 379–433.

2. Coburn, B., Sekirov, I., Finlay, B. B. (2007). Type III secretion systems and disease. Clin. Microbiol. Rev. 20, 535–549.

3. Dohlich, K., Zumsteg, A. B., Goosmann, C., Kolbe, M. (2014). A substrate-fusion protein is trapped inside the type III secretion system channel in *Shigella flexneri*. PLoS Pathog. 10, e1003881.

4. Radics, J., Königsmaier, L., Marlovits, T. C. (2014). Structure of a pathogenic type 3 secretion system in action. Nat. Struct. Mol. Biol. 21, 82–87.

5. Galan, J. E., Lara-Tejero, M., Marlovits, T. C., Wagner, S. (2014). Bacterial type III secretion systems: specialized nanomachines for protein delivery into target cells. Annu. Rev. Microbiol. 68, 415–438.

6. Deane, J. E., Abrusci, P., Johnson, S., Lea, S. M. (2010). Timing is everything: the regulation of type III secretion. Cell. Mol. Life Sci. 67, 1065–1075.

7. Barison, N., Gupta, R., Kolbe, M. (2013). A sophisticated multi-step secretion mechanism: how the type 3 secretion system is regulated. Cell. Microbiol. 15, 1809–1817.

8. Lara-Tejero, M., Kato, J., Wagner, S., Liu, X., Galan, J. E. (2011). A sorting platform determines the order of protein secretion in bacterial type III systems. Science 331, 1188–1191.

9. Morita-Ishihara, T., Ogawa, M., Sagara, H., Yoshida, M., Katayama, E., Sasakawa, C. (2006). *Shigella* Spa33 is an essential C-ring component of type III secretion machinery. J. Biol. Chem. 281, 599–607.

10. Hu, B., Lara-Tejero, M., Kong, Q., Galan, J. E., Liu, J. (2017). In situ molecular architecture of the *Salmonella* type III secretion machine. Cell 168, 1065–1074 e1010.

11. Makino, F., Shen, D., Kajimura, N., Kawamoto, A., Pissaridou, P., Oswin, H., Pain, M., Murillo, I., Namba, K., Blocker, A. J. (2016). The architecture of the cytoplasmic region of type III secretion systems. Sci. Rep. 6, 33341.

12. Diepold, A., Kudryashev, M., Delalez, N. J., Berry, R. M., Armitage, J. P. (2015). Composition, formation, and regulation of the cytosolic C-ring, a dynamic component of the type III secretion injectisome. PLoS Biol. 13, e1002039.

13. Zhang, Y., Lara-Tejero, M., Bewersdorf, J., Galan, J. E. (2017). Visualization and characterization of individual type III protein secretion machines in live bacteria. Proc. Natl. Acad. Sci. USA 114, 6098–6103.

14. Diepold, A., Sezgin, E., Huseyin, M., Mortimer, T., Eggeling, C., Armitage, J. P. (2017). A dynamic and adaptive network of cytosolic interactions governs protein export by the T3SS injectisome. Nat. Commun. 8, 15940.

15. Notti, R. Q., Bhattacharya, S., Lilic, M., Stebbins, C. E. (2015). A common assembly module in injectisome and flagellar type III secretion sorting platforms. Nat. Commun. 6, 7125.

16. Lara-Tejero, M., Qin, Z., Hu, B., Butan, C., Liu, J., Galan, J. E. (2019). Role of SpaO in the assembly of the sorting platform of a *Salmonella* type III secretion system. PLoS Pathog. 15, e1007565.

17. Song, M., Sukovich, D. J., Ciccarelli, L., Mayr, J., Fernandez-Rodriguez, J., Mirsky, E. A., Tucker, A. C., Gordon, D. B., Marlovits, T. C., Voigt, C. A. (2017). Control of type III protein secretion using a minimal genetic system. Nat. Commun. 8, 14737.

18. Yu, X. J., Liu, M., Matthews, S., Holden, D. W. (2011). Tandem translation generates a chaperone for the *Salmonella* type III secretion system protein SsaQ. J. Biol. Chem. 286, 36098–36107.

19. Bzymek, K. P., Hamaoka, B. Y., Ghosh, P. (2012). Two translation products of *Yersinia yscQ* assemble to form a complex essential to type III secretion. Biochemistry 51, 1669–1677.

20. McDowell, M. A., Marcoux, J., McVicker, G., Johnson, S., Fong, Y. H., Stevens, R., Bowman, L. A., Degiacomi, M. T., Yan, J., Wise, A., Friede, M. E., Benesch, J. L., Deane, J. E., Tang, C. M., Robinson, C. V., Lea, S. M. (2016). Characterisation of *Shigella* Spa33 and *Thermotoga* FliM/N reveals a new model for C-ring assembly in T3SS. Mol. Microbiol. 99, 749–766.

21. Lossl, P., van de Waterbeemd, M., Heck, A. J. (2016). The diverse and expanding role of mass spectrometry in structural and molecular biology. EMBO J. 35, 2634–2657.

22. Sharon, M. (2010). How far can we go with structural mass spectrometry of protein complexes? J. Am. Soc. Mass Spectrom. 21, 487–500.

23. Ngounou Wetie, A. G., Sokolowska, I., Woods, A. G., Roy, U., Loo, J. A., Darie, C. C. (2013). Investigation of stable and transient protein-protein interactions: Past, present, and future. Proteomics 13, 538–557.

24. Benesch, J. L. P. (2009). Collisional Activation of Protein Complexes: Picking Up the Pieces. J. Am. Soc. Mass Spectrom. 20, 341–348.

25. Minamino, T., MacNab, R. M. (2000). FliH, a soluble component of the type III flagellar export apparatus of *Salmonella*, forms a complex with FliI and inhibits its ATPase activity. Mol. Microbiol. 37, 1494–1503.

26. Svergun, D. I. (1999). Restoring low resolution structure of biological macromolecules from solution scattering using simulated annealing. Biophys. J. 76, 2879–2886.

27. Hu, B., Morado, D. R., Margolin, W., Rohde, J. R., Arizmendi, O., Picking, W. L., Picking, W. D., Liu, J. (2015). Visualization of the type III secretion sorting platform of *Shigella flexneri*. Proc. Natl. Acad. Sci. USA 112, 1047–1052.

28. Rocha, J. M., Richardson, C. J., Zhang, M., Darch, C. M., Cai, E., Diepold, A., Gahlmann, A. (2018). Single-molecule tracking in live *Yersinia enterocolitica* reveals distinct cytosolic complexes of injectisome subunits. Integr. Biol. 10, 502–515.

29. Case, H. B., Dickenson, N. E. (2018). MxiN differentially regulates monomeric and oligomeric species of the *Shigella* type three secretion system ATPase Spa47. Biochemistry 57, 2266–2277.

30. Datsenko, K. A., Wanner, B. L. (2000). One-step inactivation of chromosomal genes in *Escherichia coli* K-12 using PCR products. Proc. Natl. Acad. Sci. USA 97, 6640–6645.

31. Bochner, B. R., Huang, H. C., Schieven, G. L., Ames, B. N. (1980). Positive selection for loss of tetracycline resistance. J. Bacteriol. 143, 926–933.

32. Maloy, S. R., Nunn, W. D. (1981). Selection for loss of tetracycline resistance by *Escherichia coli*. J. Bacteriol. 145, 1110–1111.

33. Vidal, K., Grosjean, I., evillard, J. P., Gespach, C., Kaiserlian, D. (1993). Immortalization of mouse intestinal epithelial cells by the SV40-large T gene. Phenotypic and immune characterization of the MODE-K cell line. J. Immunol. Methods 166, 63–73.

34. Dunne, M., Leicht, S., Krichel, B., Mertens, H. D., Thompson, A., Krijgsveld, J., Svergun, D. I., Gomez-Torres, N., Garde, S., Uetrecht, C., Narbad, A., Mayer, M. J., Meijers, R. (2016). Crystal structure of the CTP1L endolysin reveals how its activity is regulated by a secondary translation product. J. Biol. Chem. 291, 4882–4893.

35. van den Heuvel, R. H., van Duijn, E., Mazon, H., Synowsky, S. A., Lorenzen, K., Versluis, C., Brouns, S. J., Langridge, D., van der Oost, J., Hoyes, J., Heck, A. J. (2006). Improving the performance of a quadrupole time-of-flight instrument for macromolecular mass spectrometry. Anal. Chem. 78, 7473–7483.

36. Morgner, N., Robinson, C. V. (2012). Massign: an assignment strategy for maximizing information from the mass spectra of heterogeneous protein assemblies. Anal. Chem. 84, 2939–2948.

37. Blanchet, C. E., Spilotros, A., Schwemmer, F., Graewert, M. A., Kikhney, A., Jeffries, C. M., Franke, D., Mark, D., Zengerle, R., Cipriani, F., Fiedler, S., Roessle, M., Svergun, D. I. (2015). Versatile sample environments and automation for biological solution X-ray scattering experiments at the P12 beamline (PETRA III, DESY). J. Appl. Crystallogr. 48, 431–443.

38. Franke, D., Petoukhov, M. V., Konarev, P. V., Panjkovich, A., Tuukkanen, A., Mertens, H. D. T., Kikhney, A. G., Hajizadeh, N. R., Franklin, J. M., Jeffries, C. M., Svergun, D. I. (2017). ATSAS 2.8: a comprehensive data analysis suite for small-angle scattering from macromolecular solutions. J. Appl. Crystallogr. 50, 1212–1225.

39. Guinier, A. (1939). La diffraction des rayons X aux très petits angles: application à l’étude de phénomènes ultramicroscopiques. Ann. Phys. (Paris) 11, 161–237.

40. Svergun, D. I. (1992). Determination of the regularization parameter in indirect-transform methods using perceptual criteria. J. Appl. Crystallogr. 25, 495–503.

41. Franke, D., Svergun, D. I. (2009). DAMMIF, a program for rapid ab-initio shape determination in small-angle scattering. J. Appl. Crystallogr. 42, 342–346.

42. Volkov, V. V., Svergun, D. I. (2003). Uniqueness of ab initio shape determination in small-angle scattering. J. Appl. Crystallogr. 36, 860–864.

43. Tuukkanen, A. T., Kleywegt, G. J., Svergun, D. I. (2016). Resolution of ab initio shapes determined from small-angle scattering. IUCrJ 3, 440–447.

44. Petoukhov, M. V., Svergun, D. I. (2015). Ambiguity assessment of small-angle scattering curves from monodisperse systems. Acta Crystallogr. D Biol. Crystallogr. 71, 1051–1058.

45. Pettersen, E. F., Goddard, T. D., Huang, C. C., Couch, G. S., Greenblatt, D. M., Meng, E. C., Ferrin, T. E. (2004). UCSF Chimera – a visualization system for exploratory research and analysis. J. Comput. Chem. 25, 1605–1612.

