## Supporting Information for "Molecular organization of soluble type III secretion system sorting platform complexes"

Figures S1-S6

Tables S1-S6

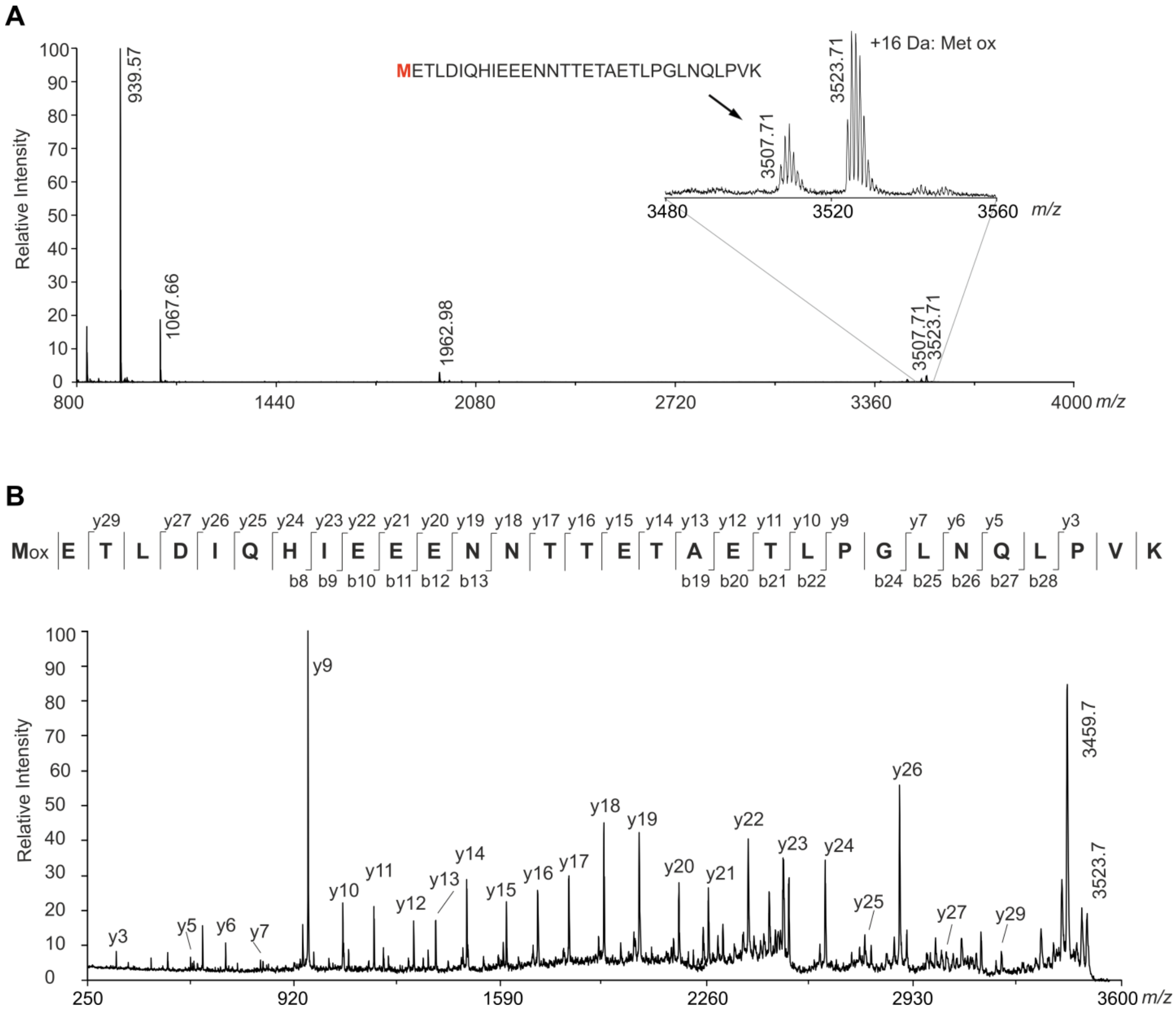


**Figure S1.** MALDI MS analysis of the N-terminal peptide of SpaO$_C$. (A) MALDI MS spectrum of the N-terminal peptide of SpaO$_C$ after in-gel tryptic digestion. (B) Amino acid sequence determination of the N-terminal peptide of SpaO$_C$. The MALDI-TOF/TOF MS/MS spectrum of the N-terminal peptide corresponding to the mass of 3523.71 Da confirms the presence of oxidized methionine in position 1. Detected y and b ions are marked by cleavage lines in the peptide sequence displayed above the spectrum. The list of all detected ions is summarized in Table S1.

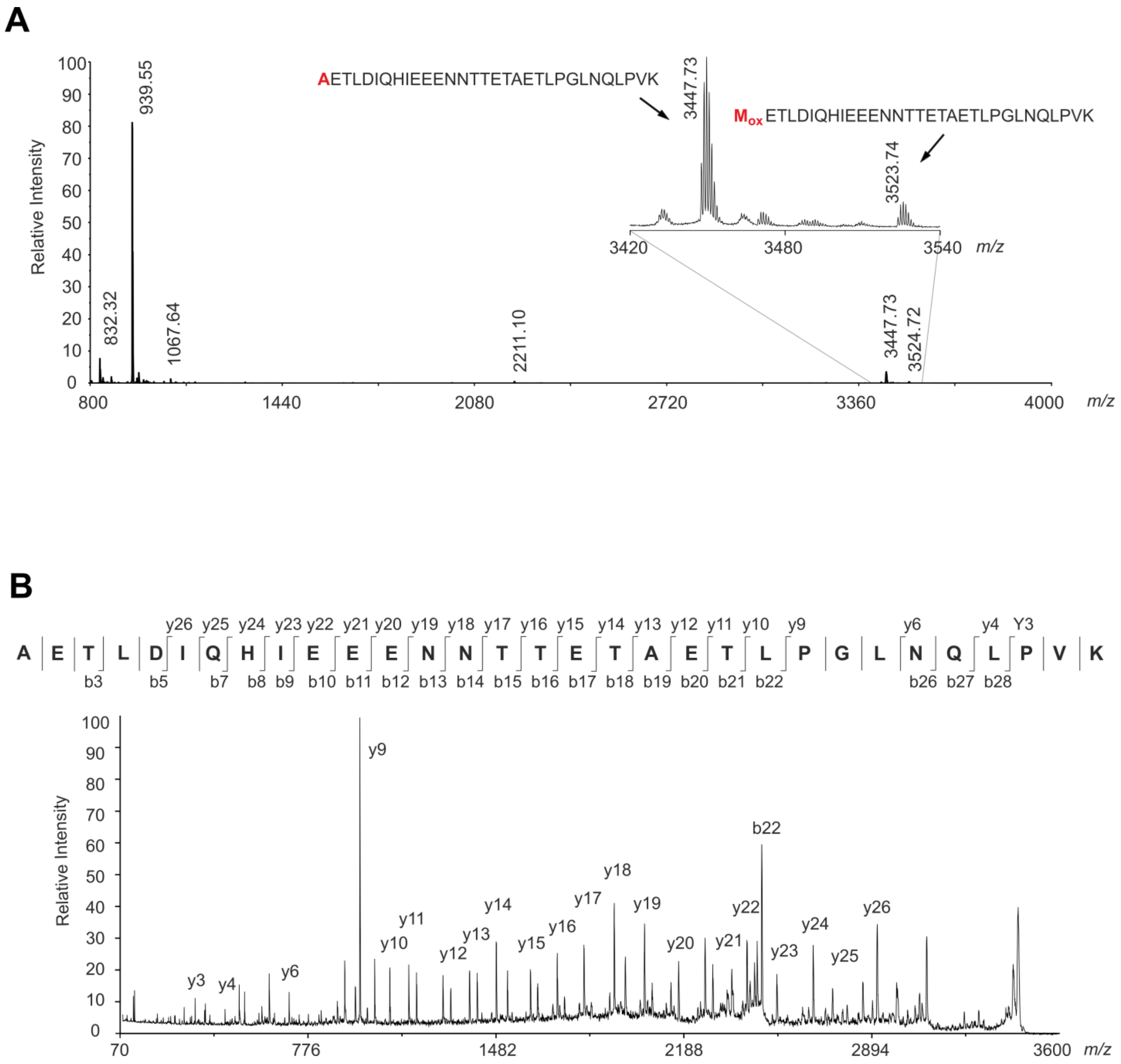


**Figure S2.** MALDI MS analysis of the N-terminal peptide of SpaO$_C$ produced by *spaO$_{V203A}$*. A) MALDI MS spectrum of the N-terminal peptide of SpaO$_C$ after in-gel tryptic digestion. (B) Amino acid sequence determination of the N-terminal peptide of SpaO$_C$. The MALDI-TOF/TOF MS/MS spectrum of the N-terminal peptide corresponding to the mass of 3447.73 Da confirming the presence of an alanine in position 1. Detected y and b ions are marked by cleavage lines in the peptide sequence displayed above the spectrum. The list of all detected ions is summarized in Table S2.

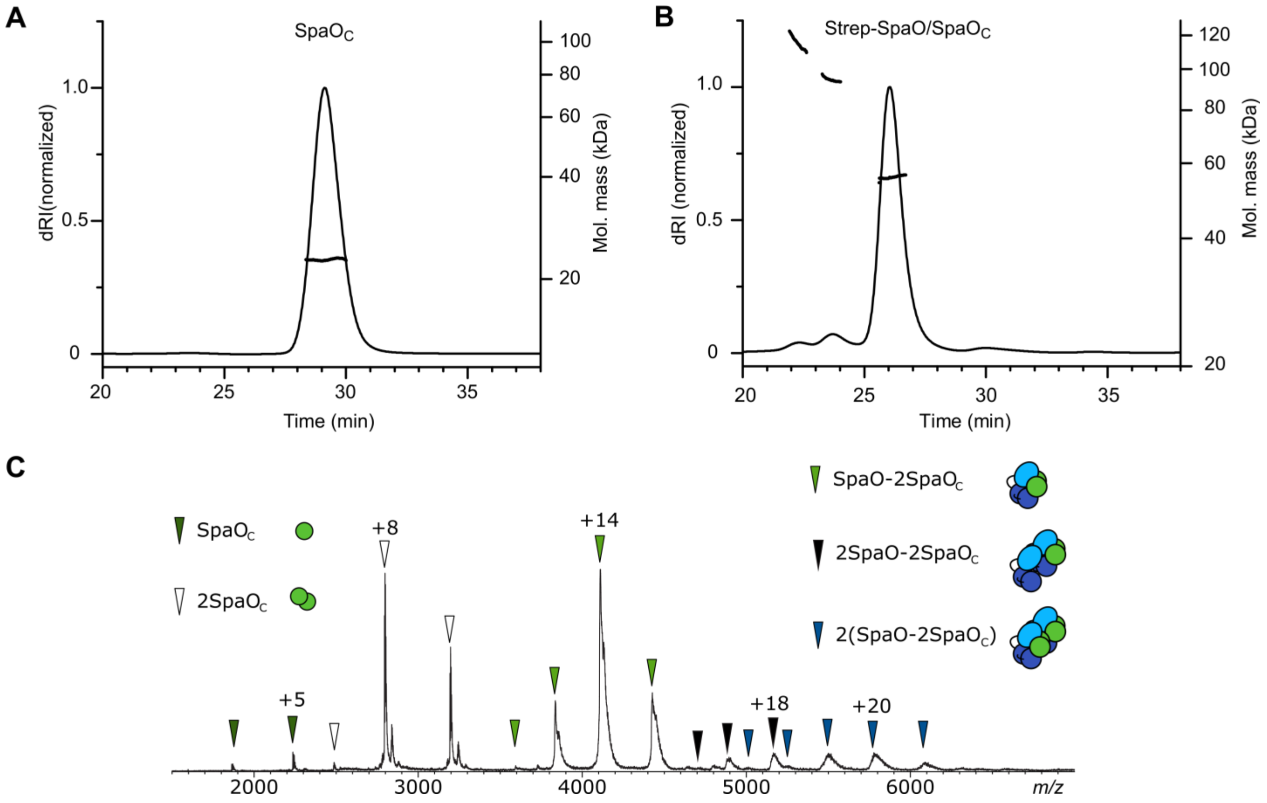


**Figure S3.** Further analysis of SpaO$_{C}$ and SpaO/SpaO$_{C}$. (A) SEC-MALS analysis of SpaO$_{C}$. The SEC elution profile (dRI trace) and the weight-averaged molar mass across the elution peak are shown. The determined mass is in good agreement with the theoretical mass of 25 kDa for dimeric SpaO$_{C}$. (B) SEC-MALS analysis of SpaO/SpaO$_{C}$ carrying an N-terminal *Strep*-tag. The experimental mass of the major species is in good agreement with the theoretical mass of 57 kDa for the SpaO-2SpaO$_{C}$ heterotrimer, while the small earlier peaks likely correspond to 2(SpaO-2SpaO$_{C}$) and 2SpaO-2SpaO$_{C}$ with theoretical masses of 115 kDa and 92 kDa, respectively. (C) Native mass spectrum of Strep-SpaO/SpaO$_{C}$. The formation of SpaO$_{C}$ dimers (white arrows), SpaO-2SpaO$_{C}$ heterotrimers (light green arrows) and dimers of SpaO-2SpaO$_{C}$ heterotrimers (blue arrows) was observed.

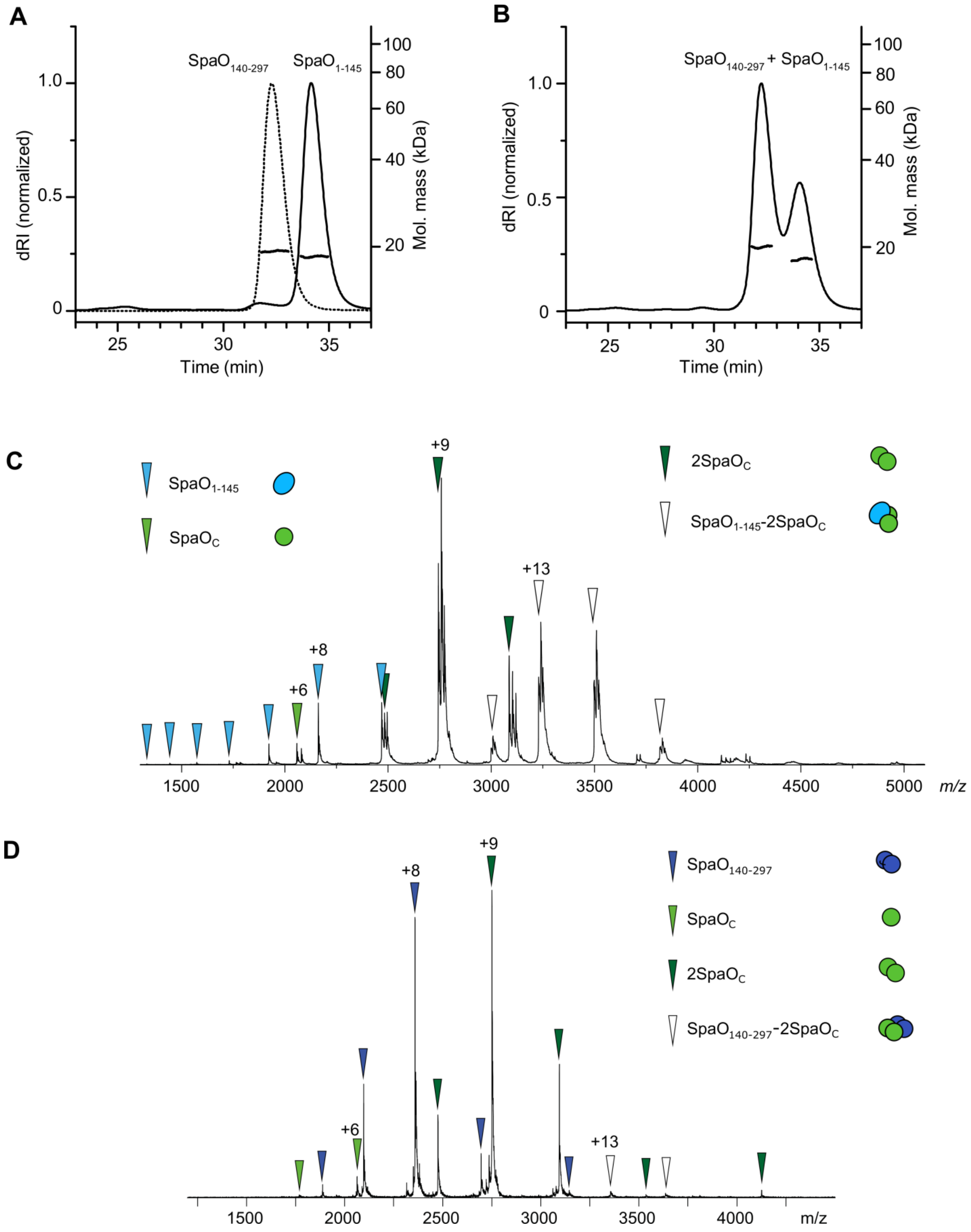


**Figure S4.** (A) SEC-MALS analysis of $SpaO_{140\text{-}297}$ (dotted line) and $SpaO_{1\text{-}145}$ (solid line). SEC elution profiles (dRI traces) and the weight-averaged molar masses across the elution peaks are shown. Experimental masses are in good agreement with the theoretical masses of monomeric $SpaO_{1\text{-}145}$ (17 kDa) and $SpaO_{140\text{-}297}$ (19 kDa). (B) SEC-MALS analysis of combined $SpaO_{1\text{-}145}$ and

$SpaO_{140-297}$. (C) Native mass spectrum of mixed $SpaO_{1-145}$ and $SpaO_C$ showing the formation of $SpaO_{1-145}$-2$SpaO_C$ complexes (white arrows). (D) Native mass spectrum of mixed $SpaO_{140-297}$ and $SpaO_C$.

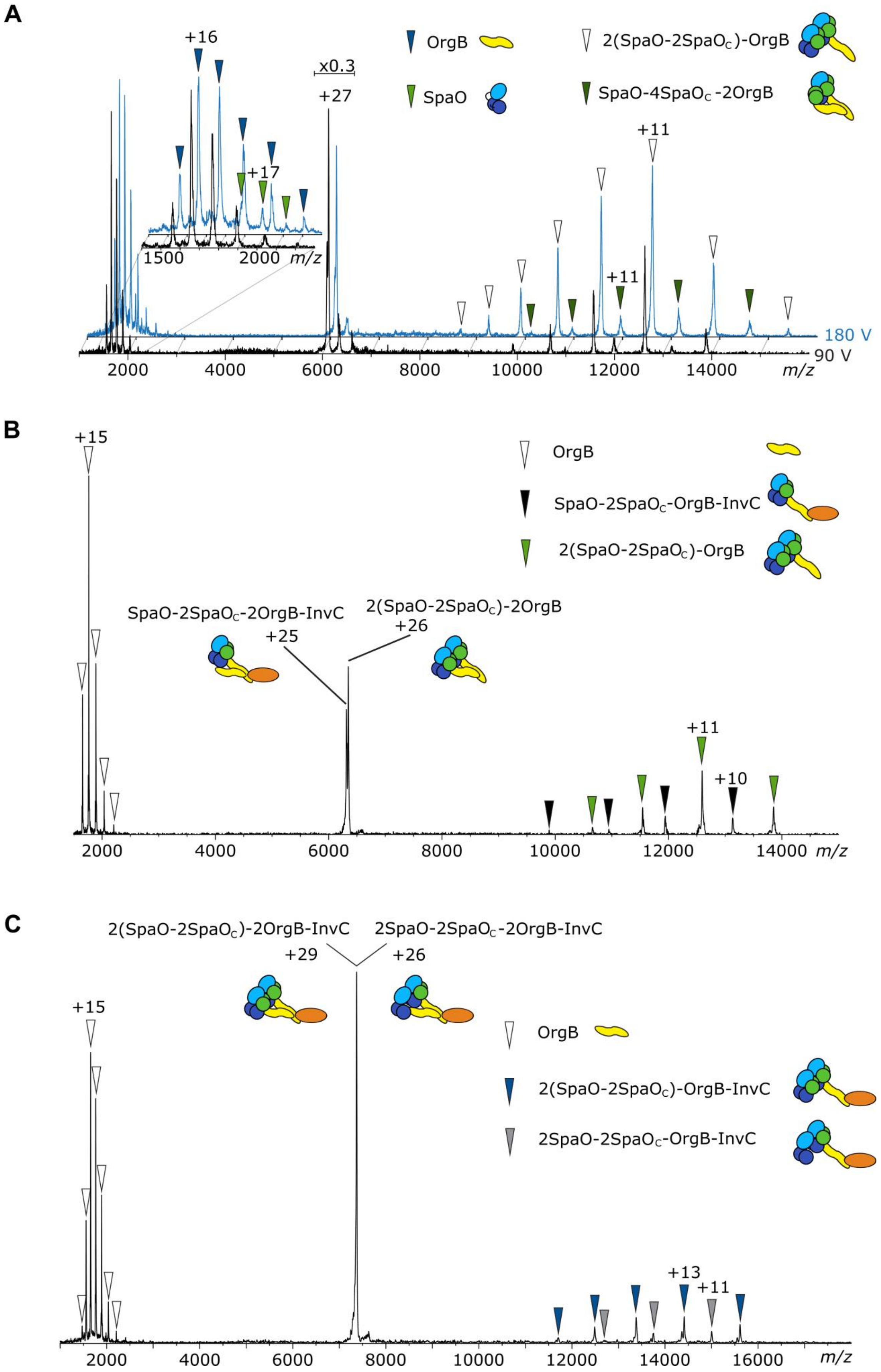
A
+16
x0.3
+27
+17
1500
2000 m/z
OrgB
SpaO
2(SpaO-2SpaOC)-OrgB
SpaO-4SpaOC -2OrgB
+11
+11
180 V
90 V
2000 4000 6000 8000 10000 12000 14000 m/z
B
+15
OrgB
SpaO-2SpaOC-OrgB-InvC
2(SpaO-2SpaOC)-OrgB
SpaO-2SpaOC-2OrgB-InvC
+25
2(SpaO-2SpaOC)-2OrgB
+26
+11
+10
2000 4000 6000 8000 10000 12000 14000 m/z
C
2(SpaO-2SpaOC)-2OrgB-InvC
2SpaO-2SpaOC-2OrgB-InvC
+29
+26
+15
OrgB
2(SpaO-2SpaOC)-OrgB-InvC
2SpaO-2SpaOC-OrgB-InvC
+13
+11
2000 4000 6000 8000 10000 12000 14000 16000 m/z

**Figure S5.** CID MS/MS analysis of sorting platform subcomplexes containing OrgB and InvC. (A) Mass spectrum of the +27 charge state of 2(SpaO-2SpaO$_C$)-2OrgB complexes (Fig. 5A) at 90 V (black spectrum) and 180 V (blue spectrum) acceleration voltage. Predominantly, OrgB (dark blue arrows) dissociates, resulting in 2(SpaO-2SpaO$_C$)-OrgB residual complexes (white arrows). To a lesser extent dissociation of SpaO (light green arrows) and residual complexes of SpaO-4SpaO$_C$-2OrgB (dark green arrows) are observed. The precursor peak in the 90 V spectrum is scaled down to 30% of its original size. (B) From the +25 charge state of SpaO-2SpaO$_C$-2OrgB-InvC complexes (Fig. 5A) an OrgB monomer (white arrows) dissociates to result in a SpaO-2SpaO$_C$-OrgB-InvC complex (black arrows). The selected peak overlapped with 2(SpaO-2SpaO$_C$)-2OrgB complexes, from which OrgB was also ejected. (C) The +26 charge state of 2SpaO-2SpaO$_C$-2OrgB-InvC and the +29 charge state of 2(SpaO-2SpaO$_C$)-2OrgB-InvC overlap (Fig. 5A) and both lose OrgB monomers, resulting in 2SpaO-2SpaO$_C$-OrgB-InvC and 2(SpaO-2SpaO$_C$)-OrgB-InvC complexes (grey and blue arrows, respectively). Experimental and theoretical molecular masses are given in Table S3.

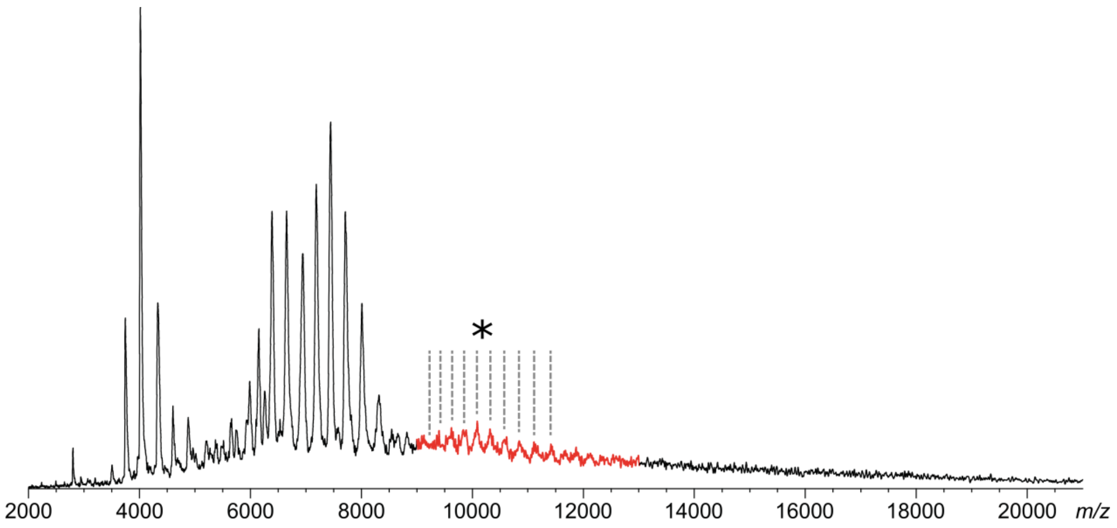


**Figure S6.** Native mass spectrum of SpaO/SpaO$_C$/OrgB/InvC complexes showing an additional peak series in the higher *m/z* range (highlighted in red). Peak positions are indicated by dashed lines.

**Table S1.** Peak list of the MALDI MS/MS spectrum of the N-terminal peptide of $SpaO_C$

| Mass | Intensity | Assigned ion | Mass | Intensity | Assigned ion |
|---|---|---|---|---|---|
| 343.18668 | 61 | y3 | 1,811.74097 | 214 | y17 |
| 382.17477 | 38 | | 1,908.74207 | 93 | |
| 456.24936 | 47 | | 1,925.74500 | 309 | y18 |
| 510.21487 | 59 | | 2,039.79761 | 235 | y19 |
| 584.29779 | 49 | y5 | 2,043.65857 | 127 | |
| 590.13623 | 36 | | 2,080.66260 | 70 | |
| 595.33911 | 40 | | 2,151.72168 | 88 | |
| 623.27216 | 115 | | 2,168.81299 | 168 | y20 |
| 698.33301 | 79 | y6 | 2,171.71826 | 100 | |
| 736.35266 | 36 | | 2,215.61000 | 43 | b19 |
| 811.37231 | 43 | y7 | 2,280.69385 | 91 | |
| 819.38165 | 40 | | 2,297.82300 | 167 | y21 |
| 831.31989 | 32 | | 2,344.67432 | 99 | b20 |
| 948.44794 | 118 | | 2,426.84839 | 217 | y22 |
| 965.45569 | 738 | y9 | 2,445.79000 | 81 | b21 |
| 984.27399 | 59 | b8 | 2,494.78516 | 141 | |
| 1,033.33374 | 46 | | 2,539.92090 | 189 | y23 |
| 1,078.52539 | 163 | y10 | 2,543.74902 | 205 | |
| 1,081.43127 | 78 | | 2,558.78101 | 203 | b22 |
| 1,097.36902 | 65 | b9 | 2,561.68604 | 116 | |
| 1,179.55261 | 156 | y11 | 2,676.90894 | 176 | y24 |
| 1,210.44080 | 69 | | 2,712.86000 | 31 | b24 |
| 1,226.40308 | 51 | b10 | 2,804.96069 | 71 | y25 |
| 1,291.44458 | 47 | | 2,825.81519 | 62 | b25 |
| 1,308.61865 | 125 | y12 | 2,918.03223 | 264 | y26 |
| 1,339.47437 | 47 | | 2,940.92212 | 97 | b26 |
| 1,355.41821 | 64 | b11 | 3,034.00098 | 82 | y27 |
| 1,379.62048 | 125 | y13 | 3,067.96000 | 45 | b27 |
| 1,480.64709 | 213 | y14 | 3,117.90283 | 85 | |
| 1,484.49255 | 75 | b12 | 3,119.01563 | 89 | |
| 1,535.44385 | 50 | | 3,180.97510 | 53 | b28 |
| 1,599.46033 | 80 | b13 | 3,231.16504 | 42 | |
| 1,609.66479 | 166 | y15 | 3,248.09351 | 61 | y29 |
| 1,710.71301 | 190 | y16 | 3,375.27344 | 82 | |

**Table S2.** Peak list of the MALDI MS/MS spectrum of the N-terminal peptide of $SpaO_C$ produced by $spaO_{V203A}$

| Mass | Intensity | Assigned ion | Mass | Intensity | Assigned ion |
|---|---|---|---|---|---|
| 110.10932 | 5,642 | | 1,711.2196 | 18,007 | y16 |
| 114.71353 | 2,122 | | 1,713.0785 | 6,224 | |
| 197.18968 | 1,258 | | 1,720.7455 | 3,189 | |
| 243.11987 | 1,246 | | 1,738.0990 | 8,656 | b15 |
| 266.16986 | 1,683 | | 1,811.2650 | 5,316 | y17 |
| 302.20706 | 2,853 | b3 | 1,812.2664 | 28,963 | |
| 343.30328 | 5,847 | y3 | 1,839.1367 | 6,980 | b16 |
| 379.31152 | 1,890 | | 1,909.2716 | 5,541 | |
| 382.30579 | 4,313 | | 1,926.3187 | 46,088 | y18 |
| 399.31384 | 1,922 | | 1,950.1892 | 5,506 | |
| 456.41013 | 1,598 | y4 | 1,953.3890 | 1,576 | |
| 483.35327 | 2,078 | | 1,968.1953 | 28,064 | b17 |
| 510.30441 | 4,808 | | 2,040.3694 | 41,723 | y19 |
| 510.38602 | 4,231 | | 2,069.2708 | 7,463 | b18 |
| 530.37787 | 6,922 | b5 | 2,123.0603 | 4,083 | |
| 584.48328 | 1,663 | | 2,140.3015 | 12,837 | b19 |
| 595.42297 | 3,426 | | 2,169.4731 | 28,567 | y20 |
| 595.51605 | 2,206 | | 2,269.3752 | 37,001 | b20 |
| 611.47723 | 1,466 | | 2,298.4932 | 25,880 | y21 |
| 619.37616 | 1,532 | | 2,353.3782 | 4,384 | |
| 623.46155 | 14,475 | | 2,370.4390 | 16,248 | b21 |
| 625.41034 | 1,806 | | 2,427.5283 | 31,269 | y22 |
| 643.49976 | 2,641 | | 2,438.4897 | 7,070 | |
| 698.53369 | 6,979 | y6 | 2,456.4099 | 11,248 | |
| 709.52020 | 1,438 | | 2,467.1709 | 6,941 | |
| 716.50146 | 1,810 | | 2,483.4988 | 80,214 | b22 |
| 736.57990 | 2,000 | | 2,540.6892 | 16,515 | y23 |
| 771.52911 | 2,267 | b7 | 2,541.5774 | 8,171 | |
| 811.65717 | 2,200 | | 2,638.4495 | 4,292 | |
| 819.56348 | 3,006 | | 2,677.7136 | 37,144 | y24 |
| 880.61121 | 3,277 | | 2,725.7891 | 2,477 | |
| 908.61090 | 20,411 | b8 | 2,749.5652 | 6,167 | |
| 912.43182 | 2,312 | | 2,750.8560 | 2,270 | |
| 947.66748 | 3,893 | | 2,752.7341 | 3,932 | |
| 948.69568 | 9,094 | | 2,805.7908 | 7,400 | y25 |
| 965.74176 | 110,188 | y9 | 2,864.6448 | 13,361 | b26 |
| 993.71252 | 2,812 | | 2,865.7361 | 7,149 | |
| 1,021.7034 | 18,715 | b9 | 2,918.9241 | 49,539 | y26 |
| 1,078.8221 | 21,098 | y10 | 2,922.7793 | 4,895 | |
| 1,150.7382 | 16,086 | b10 | 2,992.7439 | 8,700 | b27 |
| 1,179.8917 | 14,451 | y11 | 2,996.9338 | 5,841 | |
| 1,279.8339 | 16,035 | b11 | 3,081.7839 | 4,022 | |
| 1,308.9347 | 17,669 | y12 | 3,103.9639 | 3,054 | |
| 1,379.9927 | 17,100 | y13 | 3,105.9302 | 11,129 | b28 |
| 1,382.2769 | 2,175 | | 3,106.8362 | 11,304 | |
| 1,408.8663 | 18,488 | b12 | 3,107.7800 | 9,100 | |
| 1,481.0442 | 28,982 | y14 | 3,109.6184 | 3,845 | |
| 1,522.9613 | 16,351 | b13 | 3,249.0701 | 2,806 | |
| 1,610.1277 | 19,401 | y15 | 3,303.0796 | 5,021 | |
| 1,637.0116 | 17,267 | b14 | 3,304.9766 | 1,368 | |

**Table S3.** Theoretical masses and average experimental masses of proteins and protein complexes as determined by native MS (n≥3, unless otherwise stated)

| Protein/-complex | Theoretical mass (Da) | Experimental avg. mass (Da) | STDEV (Da) | Avg. FWHM (Da) |
|---|---|---|---|---|
| $SpaO_C$ MS/MS | 11,176.0 | 11,170 | 50 | 40 |
| $SpaO_C$-Strep MS/MS | 12,374.0 | 12,371 | 3 | 10 |
| $SpaO_{1-145}$-Strep | 17,291.8 | 17,292.0 | 0.9 | 12 |
| Strep-$SpaO_{140-297}$ | 18,863.6 | 18,863.7 | 0.6 | 13 |
| $2SpaO_C$ | 22,351.0 | 22,349 | 5 | 15 |
| $2SpaO_C$-Strep | 24,748.0 | 24,746.9 | 1.5 | 14 |
| OrgB MS/MS | 26,448.4 | 26,459 | 21 | 100 |
| $SpaO_{1-145}$-Strep/$SpaO_C$-Strep MS/MS | 29,665.8 | 29,640 | 11 | 130 |
| SpaO MS/MS | 33,793.7 | 33,800 | 40 | 250 |
| $SpaO_{1-145}$-Strep/$2SpaO_C$-Strep | 42,039.8 | 42,000 | 70 | 240 |
| $2SpaO_C$-Strep/Strep-$SpaO_{140-297}$ | 43,611.3 | 43,663 | 26 | 410 |
| SpaO/$SpaO_C$ MS/MS | 44,838.1 | 44,880 | 50 | 50 |
| SpaO-Strep/$SpaO_C$-Strep MS/MS | 47,524.0 | 47,504 | 23 | 70 |
| InvC-Strep | 48,808.9 | 48,240 | 110 | 370 |
| SpaO/$2SpaO_C$ | 56,013.6 | 56,050 | 40 | 190 |
| Strep-SpaO/$2SpaO_C$ | 57,471.2 | 57,552 | 19 | 270 |
| SpaO-Strep/$2SpaO_C$-Strep | 59,897.9 | 59,930 | 40 | 250 |
| $SpaO_{1-145}$-Strep/$2SpaO_C$-Strep/Strep-$SpaO_{140-297}$ | 60,903.0 | 60,980 | 50 | 460 |
| $SpaO_{1-145}$-Strep/$4SpaO_C$-Strep | 66,787.1 | 66,920 | 70 | 710 |
| 2OrgB/InvC-Strep | 101,705.6 | 102,220 | 190 | 820 |
| 2SpaO-Strep/$3SpaO_C$-Strep MS/MS | 107,422.0 | 107,390 | 50 | 190 |
| SpaO/$2SpaO_C$/2OrgB | 109,041.5 | 109,230 | 100 | 890 |
| 2(SpaO/$2SpaO_C$) | 112,027.2 | 112,480 | 170 | 530 |
| 2SpaO/$2SpaO_C$/OrgB MS/MS | 116,386.9 | 116,150 | 40 | 320 |
| 2(SpaO-Strep/$2SpaO_C$-Strep) | 119,795.7 | 120,020 | 180 | 750 |
| SpaO/$2SpaO_C$/1OrgB/InvC-Strep MS/MS | 131,402.1 | 131,350 | 90 | 240 |
| 2SpaO/$4SpaO_C$/OrgB MS/MS | 138,475.6 | 138,850 | 290 | 250 |
| 2SpaO/$2SpaO_C$/2OrgB | 142,835.2 | 143,400 | 240 | 1,300 |
| 2SpaO-Strep/$4SpaO_C$-Strep/OrgB-His MS/MS | 147,511.5 | 147,380 | 50 | 330 |
| SpaO/$2SpaO_C$/2OrgB/InvC-Strep | 157,850.4 | 158,800 | 400 | 1,300 |
| 2SpaO/$4SpaO_C$/2OrgB | 164,923.9 | 165,370 | 160 | 910 |
| 2SpaO/$2SpaO_C$/1OrgB/InvC-Strep MS/MS * | 165,195.8 | 165,019 | 28 | 260 |
| 2SpaO-Strep/$4SpaO_C$-Strep/2OrgB-His | 175,227.2 | 175,500 | 500 | 890 |
| SpaO/$4SpaO_C$/2OrgB/InvC-Strep MS/MS | 180,070.2 | 180,400 | 400 | 1,400 |
| 2SpaO/$4SpaO_C$/OrgB/InvC-Strep MS/MS | 187,284.5 | 187,620 | 170 | 490 |
| 2SpaO/$2SpaO_C$/2OrgB/InvC-Strep | 191,644.2 | 193,200 | 220 | 1,400 |
| 2SpaO/$4SpaO_C$/2OrgB/InvC-Strep | 213,732.8 | 214,500 | 500 | 1,800 |

STDEV: Standard deviation
Avg. FWHM: Average full-width at half-maximum
*n=2

**Table S4.** SAXS data acquisition, sample details, data analysis, modelling fitting and software used

| **(a) Sample details** | | | | | | |
|---|---|---|---|---|---|---|
| Organism | *Salmonella enterica*, subsp. *enterica*, serovar Typhimurium | | | | | |
| Strain | SL1344 | | | | | |
| | $SpaO_C$ | $SPAO_{140-297}$ | $SpaO_{1-145}$ | $SpaO_{1-145}$-2$SpaO_C$ | SpaO-2$SpaO_C$ | SpaO-2$SpaO_C$-2OrgB-InvC |
| UniProt ID | P40699 (203-303) | P40699 (140-297) | P40699 (1-145) | P40699 (1-145) P40699 (203-303) | P40699 (1-303) P40699 (203-303) | P40699 (1-303) P40699 (203-303) P0CL45 (1-226) E8XL22 (1-431) |
| Tags | C-term *Strep*-tag | N-term *Strep*-tag | C-term *Strep*-tag | C-term *Strep*-tag/ C-term *Strep*-tag | N-term *Strep*-tag/ no tag | N-term *Strep*-tag/ no tag/no tag/no tag |
| Extinction coefficient ε ($M^{-1}$ $cm^{-1}$, 280 nm) | 24,980 | 20,970 | 45,615 | 70,595 | 75,190 | 186,670$^{\dagger}$ & 256,110$^{\#}$ |
| Molecular mass from chemical composition (kDa) * | 25 | 19 | 17 | 42 | 58 | 158$^{\dagger}$ & 214$^{\#}$ |
| Loading/injection volume (µl) | 90 | 90 | 90 | 90 | 90 | 75 |
| Concentration, (mg $ml^{-1}$) | 14.8 | 10.2 | 6.9 | 11.1 | 8.3 | 17.6$^{\dagger}$/ 17.4$^{\#}$ |
| **SEC-SAXS** | | | | | | |
| Column type | Superdex 200 increase 10/300 GL | Superdex 200 increase 10/300 GL | Superdex 200 increase 10/300 GL | Superdex 200 increase 10/300 GL | Superdex 200 increase 10/300 GL | Superose 6 10/300 GL |
| Flow rate (ml $min^{-1}$) | 0.5 | 0.5 | 0.5 | 0.5 | 0.5 | 0.3 |
| Solvent composition and source | 20 mM HEPES pH 7.5, 150 mM NaCl (in water) | 20 mM HEPES pH 7.5, 150 mM NaCl (in water) | 20 mM HEPES pH 7.5, 150 mM NaCl (in water) | 20 mM HEPES pH 7.5, 150 mM NaCl (in water) | 20 mM HEPES pH 7.5, 150 mM NaCl (in water) | 10 mM Tris-HCL pH 8.0, 50 mM NaCl (in water) |

| **(b) SAS data collection parameters** | |
|---|---|
| Source, instrument and description or reference | P12 (EMBL/DESY, storage ring PETRA III, Germany) |
| Wavelength (Å) | 1.24 |
| Beam geometry (size, sample-to-detector distance) | 0.2 x 0.12 mm$^2$, 3.0 m |
| *q*-measurement range (Å$^{-1}$) | 0.008 – 0.47 |
| Method for monitoring radiation damage, X-ray dose | BECQUEREL software |
| Exposure time, number of exposures | 1 s, 3600x |
| Sample configuration: flow rate | 0.3 ml/min for SpaO/SpaO$_C$/OrgB/InvC, all others 0.25 ml/min |
| Sample temperature (K) | 283 |

| **(c) Software employed for SAS data reduction, analysis and interpretation** | |
|---|---|
| SAS data reduction to sample–solvent scattering, and extrapolation, merging, desmearing | PRIMUS |
| Calculation of ε from sequence | PROTPARAM |
| Basic analyses: Guinier, *P*(*r*), scattering particle volume (*e.g.* Porod volume $V_P$ or volume of correlation $V_c$) | PRIMUS |
| Shape/bead modelling | DAMMIF, SASRES |
| Molecular graphics | PYMOL |

| **(d) Structural parameters** | | | | | | |
|---|---|---|---|---|---|---|
| | SpaO$_C$ | SPAO$_{140-297}$ | SpaO$_{1-145}$ | SpaO$_{1-145}$-2SpaO$_C$ | SpaO-2SpaO$_C$ | SpaO-2SpaO$_C$-2OrgB-InvC |
| **Guinier Analysis** | | | | | | |
| I(0) (Arbitrary units) | 4,561.91 ± 8.77 | 2,968.16 ± 7.64 | 1,173.37 ± 7.9 | 4,750.57 ± 10.0 | 4,373.58 ± 14.2 | 4,149.32 ± 71.83 |
| $R_g$ (nm) | 2.5 ± 0.3 | 2.1 ± 0.2 | 1.6 ± 0.2 | 3.1 ± 0.3 | 3.3 ± 0.3 | 5.7 ± 0.6 |
| *q*-range (Å$^{-1}$) | 0.0229 - 0.3 | 0.01769 - 0.3 | 0.03006 - 0.3 | 0.01973 - 0.3 | 0.014678 - 0.3 | 0.01813 - 0.3 |
| Fidelity of Primus Guinier analysis | 0.96 | 0.93 | 0.66 | 0.91 | 0.92 | 0.63 |
| | SpaO$_C$ | SPAO$_{140-297}$ | SpaO$_{1-145}$ | SpaO$_{1-145}$-2SpaO$_C$ | SpaO-2SpaO$_C$ | SpaO-2SpaO$_C$-2OrgB-InvC |
| ***P*(*r*) analysis** | | | | | | |

| | | | | | | |
|---|---|---|---|---|---|---|
| I(0) (arbitrary units) | 4,663.00 ± 466 | 2,937.00 ± 294 | 1,206.00 ± 121 | 4,824.00 ± 482 | 4,404.00 ± 440 | 4,592.00 ± 459 |
| $R_g$ (nm) | 2.7 ± 0.3 | 2.1 ± 0.2 | 1.7 ± 0.2 | 3.1 ± 0.3 | 3.3 ± 0.3 | 7.0 ± 0.7 |
| $D_{max}$ (nm) | 9.1 ± 0.9 | 7.2 ± 0.7 | 5.3 ± 0.5 | 10.6 ± 1.1 | 11.1 ± 1.1 | 22.7 ± 2.3 |
| *q*-range (Å$^{-1}$) | 0.0229 - 0.3 | 0.01769 - 0.3 | 0.03006 - 0.3 | 0.01973 - 0.3 | 0.014678 - 0.3 | 0.01813 - 0.3 |
| Volume Porod (nm$^3$) | 51 | 35 | 31 | 68 | 108 | 302 |
| **(e) Shape modelling results (DAMMIF)** | | | | | | |
| *q*-range for fitting (Å$^{-1}$) | 0.0229 - 0.3 | 0.01769 - 0.3 | 0.03006 - 0.3 | 0.01973 - 0.3 | 0.014678 - 0.3 | 0.01813 - 0.3 |
| Ambiguity measured by AMBIMETER | 2.494 | 0.6990 | 0.4771 | 2.509 | 2.526 | 2.127 |
| SASRES resolution (Å) | 31 ± 3 | 21 ± 2 | 31 ± 2 | 34 ± 3 | 41 ± 3 | 64 ± 5 |
| MW estimate (kDa) | 34 | 20 | 16 | 47 | 52 | 208 |
| **(f) SASBDB data and model deposition IDs** | SASDC68 | SASDEK7 | SASDC88 | SASDC98 | SASDC78 | SASDEJ7 |

* based on stoichiometry from native MS/MALS.

† based on SpaO-2SpaO$_C$-2OrgB-InvC stoichiometry

### based on 2SpaO-4SpaO$_C$-2OrgB-InvC stoichiometry

**Table S5.** Oligonucleotides used for cloning and mutagenesis of *Salmonella* genes

| **Construct** | **Oligonucleotide Sequence 5´ - 3´** | |
|---|---|---|
| *S.* Typhimurium Δ*spaO* | Fw | GCTATTGGCGCAAACCGCGACAGAATGCCAGCGCCATGGCCGGGAAGCGAGTGTAGGCTGGA<br>GCTGCTTCGA |
| | Rv | CGCCTAAGGTGTCATTCATCTGTACCAGTTCGCCATTACCCAGCAAAACAATGGGAATTAGCC<br>ATGGTCCAT |
| *S.* Typhimurium Δ*spaO*$_{C}$ | Fw 1 | TTATTGCTACGCGAAAAAGTTAGGTCATTTCAACCGTGTTTTAAGACCCACTTTCACATT |
| | Rv 1 | TATTATTTTCTTCTTCGATATGTTGAATATCTAACGTTTC CTAAGCACTTGTCTCCTG |
| | Fw 2 | TTATTGCTACGCGAAAAAGTTAGGTCATTTCAACCGTGTTGAGGGCGGCATTATTGTTGAAAC<br>GTTAGATATTCAACATATCGAA |
| | Rv 2 | CACCATTCGCCATAATTTCA |
| *S.* Typhimurium Δ*spaO*$_{FL}$ | Fw 1 | AGGATGACGCCTGATGTCATTGCGTGTGAGACAGATTGATTTAAGACCCACTTTCACATT |
| | Rv 1 | GGCGCTGGCATTCTGTCGCGGTTTGCGCCAATAGCCATTCCTAAGCACTTGTCTCCTG |
| | Fw 2 | AGGATGACGCCTGATGTCATTGCGTGTGAGACAGATTGATTGATGAGAATGGCTATTGGCGCA<br>AAC |
| | Rv 2 | ACTCAAACGGTCGCTCTGTT |
| *S.* Typhimurium *spaO-3XFLAG*<br>*S.* Typhimurium Δ*spaO*$_{C}$*-3XFLAG*<br>*S.* Typhimurium Δ*spaO*$_{FL}$*-3XFLAG* | Fw 1 | TGAGATCCATGAATGGCTGAGCGAGTCTGGTAATGGGGAATTAAGACCCACTTTCACATT |
| | Rv 1 | CAGGGTGGAAAATGCCAGTAAGGCAATTAATGAGATATCACTAAGCACTTGTCTCCTG |
| | Fw 2 | TGAGATCCATGAATGGCTGAGCGAGTCTGGTAATGGGGAACGGGCTGACTACAAAGACCATG<br>ACGGTGATTATAAAGATCATGA<br>CATCGACTACAAGGATGACGATGACAAGTAGTGAGG |
| | Rv 2 | AACGAAACAGGTTCCTGACGCAATAATAAATGGCAACAGGGTGGAAAATGCCAGTAAGGCAA<br>TTAATGAGATATCATTCCCCAT<br>TACCAGACTCGCTCAGCCTCACTACTTGTCATCGTC |
| pASK-IBA3+ *spaO*$_{C}$*-Strep* | Fw | AAAAAAGGTCTCAAATGGAAACGTTAGATATTCA |
| | Rv | AAAAAAGGTCTCGGCGCTTTCCCCATTACCAGACTCGC |
| pASK-IBA5+ *spaO-Strep* | Fw | TTTTTGCTAGCATGTCATTGCGTGTGAGAC |
| | Rv | CAGTCGACCTCGAGTTATCATTTTTCAAACTGCGGATGGCTCCACGCGCTTTCCCCATTACCAG<br>ACTCGC |
| pASK-IBA5+ *Strep-spaO* | Fw | AAAAAAGGTCTCGGCGCCATGTCATTGCGTGTGAGACA |
| | Rv | AAAAAAGGTCTCGTATCATTCCCCATTACCAG |
| pASK-IBA5+ *spaO*$_{V203A}$*-Strep* | Fw | GGGGGAATTATTGCGGAAACGTTAGAT |
| | Rv | ATCTAACGTTTCCGCAATAATTCCCCC |
| pASK-IBA3C *spaO*$_{1-145}$*-Strep* | Fw | ACGTGTGGTCTCGAATGTCATTGCGTGTGAGACA |
| | Rv | ACGTGTGGTCTCCGCGCTCGGCCTGCCGCCCCCGACTG |
| pASK-IBA5+ *Strep-spaO*$_{140-297}$ | Fw | ACGTGTGGTCTCCGCGCCCGGGGGCGGCAGGCCGAAAA |
| | Rv | ACGTGTGGTCTCGTATCAGCTCAGCCATTCATGGATCT |

| | | |
|---|---|---|
| pET28a *orgB*-His | Fw<br>Rv | ACATGCCATGGGTATGCTCAAAAATATCCCAATACC<br>AAACCGCTCGAGCCTTATAACCTCCGCTTGCG |
| pCDF-Duet1 *orgB+spaO* *(spaO* cloning*)* | Fw<br>Rv | AATTAACATATGTCATTGCGTGTGAGACAG<br>AATTAAGGTACCTCATCATTCCCCATTACCAGACTCG |
| pCDF-Duet1 *orgB+spaO* *(orgB* cloning*)* | Fw<br>Rv | AATTAACCATGGTCAAAAATATCCCAATACCGTCC<br>AATTAAGGATCCTCATCACCTTATAACCTCCGCTTGCG |
| pCDF-Duet1 *orgB*-His | Fw<br>Rv | AATTAACCATGGTCAAAAATATCCCAATACCGTCC<br>AATTAAGGATCCTCATCAGTGGTGGTGGTGGTG |
| pACYC-Duet1 *invC-Strep* | Fw<br>Rv | AATTAACCATGGGTATGAAAACACCTCGTTTACTGCAATATC<br>AATTAAAAGCTTTTATTATTTTTCGAACTGCGGGTGG |
| pASK-IBA3+ *invC-Strep* | Fw<br>Rv | AATTAAGGTCTCTAATGAAAACACCTCGTTTACTGCAATATC<br>AATTAAGGTCTCTGCGCTATTCTGGTCAGCGAATGCATTC |
| pET28a *orgB*$_{1\text{-}105}$-His | Fw<br>Rv | AATTAACCATGGGT ATGCTCAAAAATATCCCAATACC<br>AATTAACTCGAGGACCGCAGCTGAAAATAACTC |
| pET28a *orgB*$_{106\text{-}226}$-His | Fw<br>Rv | AATTAACATATGGACCATCCCGAAACGCTTTTAAC<br>AATTAACTCGAGTCATCACCTTATAACCTCCGCTTGCG |
| pASK-IBA3C+RBS* *invC*$_{1\text{-}79}$-*Strep* | Fw<br>Rv | AATTAAGGTCTCTAATGAAAACACCTCGTTTACTGCAATATC<br>AATTAAGGTCTCTGCGCTAGTGGGATAAAGCACGACATC |
| pASK-IBA3+ *invC*$_{80\text{-}431}$-*Strep* | Fw<br>Rv | AATTAAGGTCTCTAATGGGACGTGCGTTATCGGCGTG<br>AATTAAGGTCTCTGCGCTATTCTGGTCAGCGAATGCATTC |

* pASK-IBA3C+RBS is a variant of pASK-IBA3C in which the RBS has been replaced by that of pASK-IBA3+.

**Table S6.** Constructs used for recombinant gene expression

| **Purification for biophysical analysis** | |
|---|---|
| **Protein complex** | **Constructs (co-)transformed** |
| SpaO$_{c}$ | pASK-IBA3+ *spaO$_{C}$-Strep* |
| SpaO$_{1\text{-}145}$ | pASK-IBA3C *spaO$_{1\text{-}145}$-Strep* |
| SpaO$_{1\text{-}145}$/SpaO$_{C}$ | pASK-IBA3C *spaO$_{1\text{-}145}$-Strep*, pASK-IBA3+ *spaO$_{C}$-Strep* |
| SpaO$_{140\text{-}297}$ | pASK-IBA5+ *Strep- spaO$_{140\text{-}297}$* |
| SpaO/SpaO$_{C}$ | pASK-IBA5+ *spaO-Strep* or pASK-IBA5+ *Strep-spaO* |
| SpaO/SpaO$_{C}$/OrgB | pASK-IBA5+ *spaO-Strep*, pET28a *orgB*-His |
| SpaO/SpaO$_{C}$/OrgB/InvC | pCDF-Duet1 *orgB+spaO*, pASK-IBA3+ *invC-Strep* |
| OrgB/InvC | pASK-IBA3C+RBS* *invC-Strep*, pET28a *orgB* |

| **Effect of SpaO$_{C}$ on solubility of mutant SpaO** | |
|---|---|
| **SpaO variant** | **Construct** |
| SpaO | pASK-IBA5+ *spaO-Strep* |
| SpaO$_{V203A}$ | pASK-IBA5+ *spaO$_{V203A}$-Strep* |
| SpaO$_{V203A}$ + SpaO$_{C}$ | pASK-IBA5+ *spaO$_{V203A}$-Strep*, pASK-IBA3C *spaO$_{C}$-Strep* |

| **Co-purifications of OrgB and InvC fragments** | |
|---|---|
| **Protein combination** | **Constructs (co-)transformed** |
| OrgB$_{1\text{-}105}$ | pET28a *orgB$_{1\text{-}105}$*-His |
| OrgB$_{1\text{-}105}$ + InvC | pET28a *orgB$_{1\text{-}105}$*-His, pASK-IBA3+ *invC-Strep* |
| OrgB$_{1\text{-}105}$ + SpaO/SpaO$_{C}$ | pET28a *orgB$_{1\text{-}105}$*-His, pASK-IBA5+ *spaO-Strep* |
| OrgB$_{106\text{-}226}$ | pET28a His-*orgB$_{106\text{-}226}$* |
| OrgB$_{106\text{-}226}$+ InvC | pET28a His-*orgB$_{106\text{-}226}$,* pASK-IBA3+ *invC-Strep* |
| OrgB$_{106\text{-}226}$+ SpaO/SpaO$_{C}$ | pET28a His-*orgB$_{106\text{-}226}$,* pASK-IBA5+ *spaO-Strep* |
| InvC$_{1\text{-}79}$ + OrgB | pASK-IBA3C+RBS* *invC$_{1\text{-}79}$-Strep,* pCDFDuet-1 *orgB*-His |
| InvC$_{80\text{-}431}$+ OrgB | pASK-IBA3+ *invC$_{80\text{–}431}$-Strep*, pCDFDuet-1 *orgB*-His |
| InvC + OrgB | pACYCDuet-1 *invC-Strep*, pCDFDuet-1 *orgB*-His |
| OrgB | pCDFDuet-1 *orgB*-His |

* pASK-IBA3C+RBS is a variant of pASK-IBA3C in which the RBS has been replaced by that of pASK-IBA3+.
